# Predictability of the antigenic evolution of human influenza A H3 viruses

**DOI:** 10.1101/770446

**Authors:** R. Aguas, N.M. Ferguson

## Abstract

The current influenza A antigenic evolution paradigm suggesting that antigenic evolution is highly constrained, with successful new viruses being near optimal at maximizing their antigenic distance from past strains. This begs the question of whether influenza’s antigenic evolution is fundamentally predictable, or if it takes place on a much higher dimensional antigenic space with multiple possible trajectories. We tackle this issue by building a genotype to phenotype map validated on historical hemagglutination inhibition assay data by using machine learning methods. This map uses amino acid physiochemical properties for inference, learning the expected antigenic distance given the differences in polarity and hydrophobicity observed across any two viral sequences, and is thus applicable to newly sampled viruses with previously unseen amino acids. This allows us to accurately blindly predict the antigenic relevance of soon to be vaccine viral strains. We couple the genotype to phenotype map with a molecular evolutionary simulation algorithm to explore the limits of influenza’s antigenic evolution and infer to what extent it is in fact predictable. Although we do uncover some canalization of antigenic trajectories, we find that multiple antigenic lineages are equally viable at any one point in time even though typically only one of those trajectories is actually realized.

## Introduction

Seasonal influenza epidemics remain a major public health concern, resulting in 250,000 to 500,000 deaths per year [1]. Human influenza A evolution is characterized by rapid evolution yet limited extant genetic diversity at any point in time – resulting in a characteristic single trunk phylogeny [2, 3]. At the molecular level, antigenic drift is mainly determined by mutations in two viral segments encoding for surface proteins hemagglutinin (HA) and neuraminidase (NA), which are in direct contact with the host’s immune system [4, 5]. Hemagglutination inhibition (HI) assays provide an empirical representation of antigenic drift by measuring differential antibody neutralizing responses across a panel of viruses. Antigenic maps built from HI data reveal a punctuated evolutionary pattern, with periods of relative stasis (defining antigenic clusters) followed by selective sweeps by antigenically novel strains (cluster jumps)[3]. The linearity of these antigenic maps suggests that antigenic evolution is highly constrained, with successful new viruses being near optimal at maximizing their antigenic distance from past strains. One crucial standing question is then the extent to which antigenic evolution is fundamentally predictable, or if antigenic evolution takes place on a much higher dimensional antigenic space with multiple possible trajectories. Here we address these questions using machine learning methods to predict antigenic distances from differences in physiochemical properties in influenza HA1 sequences. The genotype to phenotype (GP) map thus formulated for influenza antigenicity can reproduce the observed features of the antigenic evolution of influenza A/H3N2 viruses and accurately predicts the antigenic novelty of viruses one year ahead. By coupling our GP map with a molecular evolution algorithm, we unveil how HA1 as a whole has evolved near neutrally whilst epitope sites have undergone strong positive selection and give insights into the predictability of flu antigenic evolution. Our results suggest that influenza‘s observed evolutionary pathway is but one of many plausible trajectories, with mutations in antigenically relevant sites generating a breadth of antigenically novel viruses later subject to epidemiological selection pressures that limit the observed viral diversity.

Modelling studies have generated competing hypotheses about the key evolutionary and epidemiological determinants of observed influenza A evolutionary patterns [6–9], but definitive understanding is lacking. Influenza A phylodynamics have been simulated assuming comparably parsimonious, yet fundamentally different mechanistic processes: short-term subtype transcending immunity [6]; positive selection associated with cluster transitions in an otherwise neutrally evolving network [7]; reuse of a limited number of antigenic combinations [8]; accumulation of deleterious mutations genetically linked to beneficial ones. More recently, a more phenomenological model has successfully reproduced observed trends in influenza evolution and epidemiology [10].

One important question where existing models differ is the extent to which antigenic evolution is fundamentally predictable [10, 11] – or even deterministic [10] – or if the space of possible trajectories is much higher dimensional, with but one realized trajectory (e.g. due to stochasticity [7] or pruning of alternative lineages [6]). In a more narrow sense, recent studies have put forward algorithmic predictions of which genetic variants of an emerging viral set [12, 13] will come to dominate circulating viruses in the next year or two (and thus be suitable vaccine candidate strains). However, predicting the antigenic properties of previously unseen strains multiple years ahead remains an open challenge. Whether or not it’s possible to predict *de novo* which strains will emerge in the future ahead of their identification at even low frequency is completely unclear; the answer fundamentally depends on the extent to which influenza evolution is deterministic.

Here we address these questions using machine learning methods to predict the expected antigenic distance between two influenza A/H3N2 viruses given the differences in physiochemical properties across their respective HA1 sequences. We focused on HA1 since it appears particularly relevant for viral antigenic evolution, being the principal target of antibody mediated immunity [14, 15] and containing several empirically established antigenic sites [16, 17].

## Results

### Genotype to Phenotype Map

The HI assay outputs a viral titer which indicates the number of times the sera needed to be diluted for hemagglutination of the antigen and red blood cells to occur [18]. This suggests that there are electrostatic interactions between antigen and antibodies which are disrupted when the serum is too diluted. We thus postulated that physiochemical property differences between the amino acid sequences of a given antigen/serum pair can serve as a proxy for the antigenic distances observed in HI assays, thus formalizing a genotype-to-phenotype (GP) map. To do so, we implemented a machine learning algorithm that was informed on how differences in 54 distinct physiochemical properties relate to empirical HI titre measurements – more detailed description of this method can be found in S1 Text). Out of those 54 properties, a subset of 7 (*α* helix, *β* sheet, *β* turn, bulkiness, coil, polarity and hydrophobicity) emerged as the most relevant in explaining the observed phenotype, with polarity and hydrophobicity being the most significant. S1 Figure clearly shows that differences in hydrophobicity and polarity found in HA1 have a very high degree of concordance with genetic distances across influenza H3 viral sequences collected over time as well as historic antigenic maps [3]. By using vectors of amino acid hydrophobicity and polarity rather than the actual amino acid sequences, we allow the resulting GP map to be used to predict the antigenic characteristics of novel influenza strains with previously unseen amino acid polymorphisms. We constructed the GP map using historic HI data [3] and corresponding HA1 sequences (S1 Table), by training a Random Forest Algorithm (RFA) to learn the expected antigenic distances between two viruses (measured in integer 2-fold dilutions given by the HI assays) given the differences in amino acid polarity and hydrophobicity across their respective viral sequences. The resulting classifier is able to predict the measured antigenic distances exactly for ∼90% of the antigen/serum pairs (Figure 1A) and successfully reproduces the antigenic map of the H3N2 subtype (Figure 1B).

**Figure 1.**
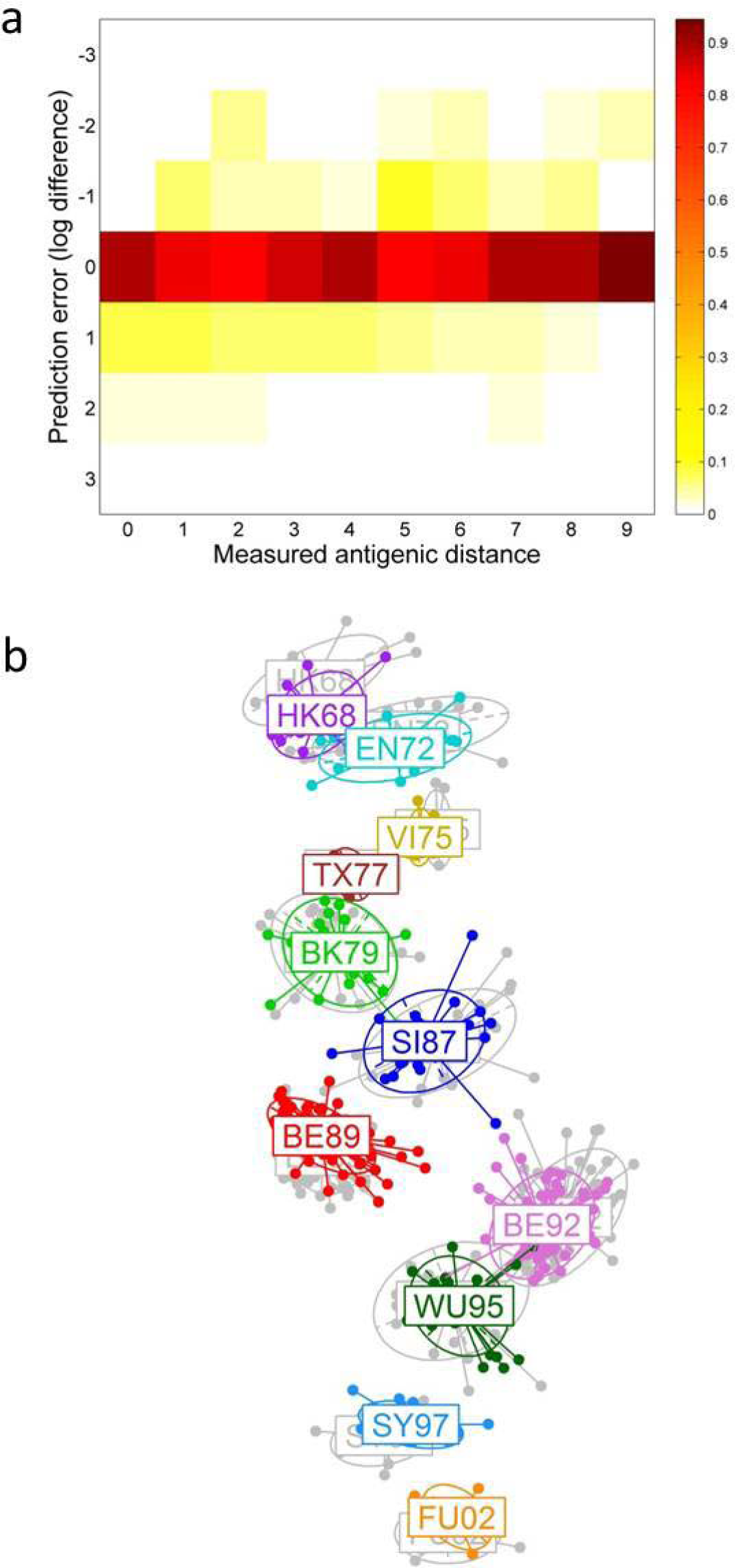
Classification accuracy of the RFA using hydrophobicity and polarity as antigenic proxy variables. (A) Surface plot showing the error in prediction of antigenic distance (distance measured on a log-2 scale) for every antigen-serum pair. Antigen-serum pairs are grouped by measured antigenic distance on the horizontal axis. The colour of each rectangular pixel illustrates the distribution of predictive errors for pairs with a particular measured antigenic. A measure of 90% for prediction error 0 indicates that 90% of antigen/serum pairs antigenic distances are predicted exactly. (B) Superimposed 2-dimensional antigenic maps generated from the RFA predicted antigenic distances (in colour) and from the HI titre data (in grey). See Methods for details of the maximum-likelihood algorithm used to construct these maps.

### Predicting the antigenic novelty of previously unseen viruses

While Figure 1 demonstrates the ability of the GP map to reproduce the data on which it was trained, this approach is also able to predict the antigenic characteristics of strains not used to train the RFA. To determine whether newly sampled viruses are antigenically novel, we train the RFA up to a specific point in time and predict the antigenic distance between all antigen/serum pairs constituted by an element of the current antigenic cluster and another element collected in the year following the reference time point. Mean one year ahead antigenic distances predicted for each year of collection show a remarkable agreement with the measured antigenic distances, particularly highlighting the antigenic novelty of future vaccine strains – Figure 2. These mean values are a compound of predicted distances to future members of the current antigenic cluster and to members of new yet unobserved clusters, since some viruses collected in cluster transition years such as 2002 e.g., can still be members of the currently circulating cluster at the time of prediction (SY97 in this example) - S2 Table. S2 Figure further illustrates the RFA prediction errors resulting from out-of-sample predictions, particularly across clusters. Given a GP map trained up to data collected in 1995, predicted antigenic distances for viruses appearing one year into the future are very similar to the measured distances (>75% having an error of ±1 log-2 unit of antigenic distance or less), as might be expected given antigenic variation is quite limited within the same antigenic cluster (for reference, the mean antigenic distance between viruses in adjacent clusters is 4.46 (±1.3)). The following year saw a cluster transition (WU95→SY97), which reduced predictive accuracy somewhat since cluster transitions are typically associated with the fixation of previously unrecorded alleles [11, 19], meaning the RFA is potentially blinded to the significance of mutations in those amino acids. Hence, while just under 70% of predicted distances are within 1 log-2 unit of the observed antigenic distance when looking 2 to 8 years into the future, a small proportion of observations have higher prediction errors associated with them, and this proportion increases slowly the further ahead one tries to predict. However, our GP map can predict the qualitative pattern of antigenic evolution even across two antigenic cluster jumps (S2C Figure).

**Figure 2.**
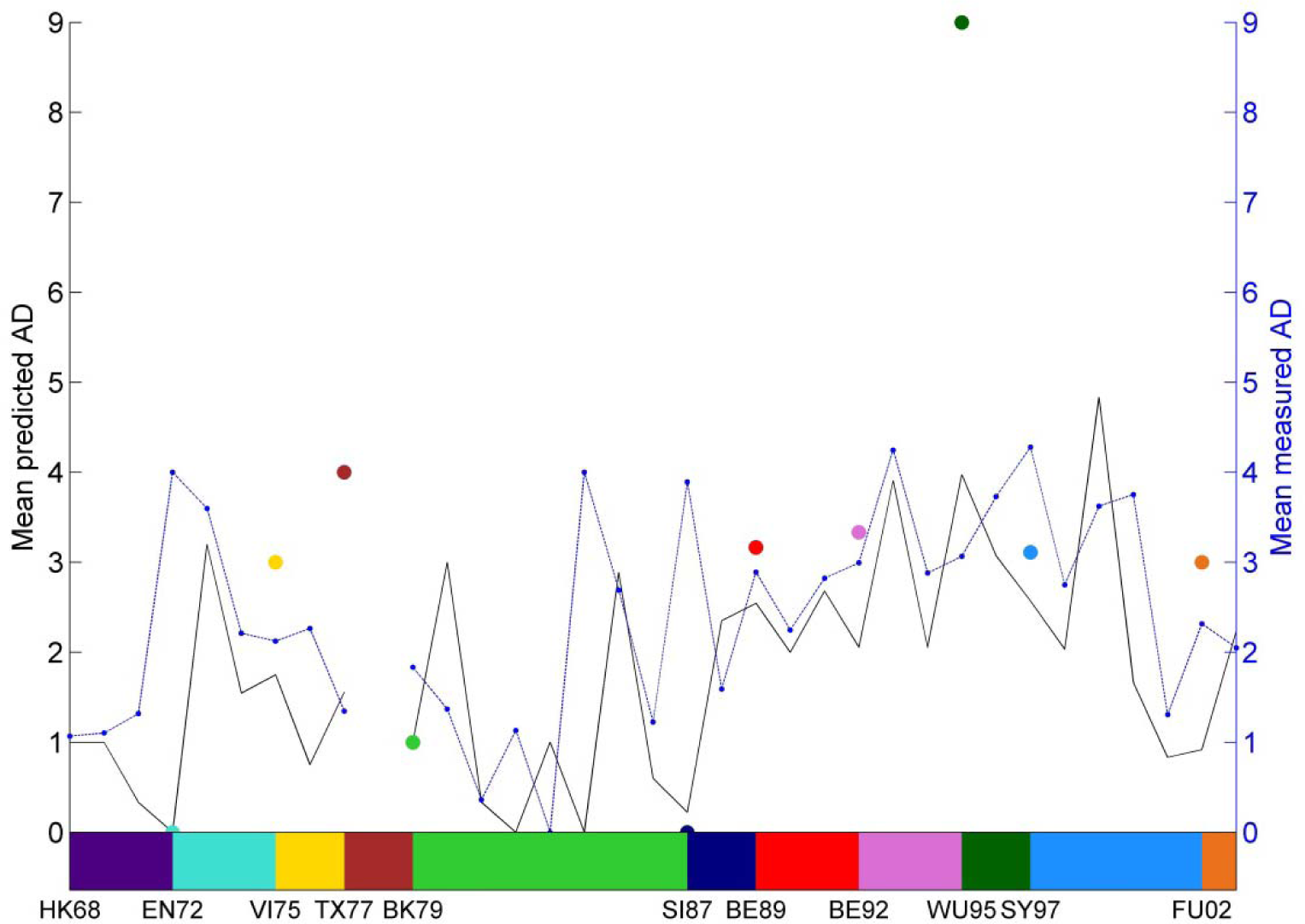
Predicting the antigenic novelty of newly sequenced viruses one year ahead. The black line shows the mean predicted antigenic distance between antigens collected in each year and sera collected in the three previous years, given an RFA trained on HI data containing antigen/serum pairs sampled up to the previous year. The colored dots identify the mean predicted antigenic distances referring to antigens which turned out to be vaccine strains, whereas the blue line represents the mean empirical antigenic distances observed for all antigen/serum pairs used for prediction each year. The different colors map vaccine strains onto their antigenic cluster, longitudinally represented as colored blocks.

### Influenza’s molecular evolution and selection for differences in physiochemical properties

The biological plausibility of our approach can be investigated by comparing the amino-acid positions in HA1 which the RFA identifies as significant in predicting antigenicity, with those of known epitopes (Figure 3). We find that the set of significant positions identified by the RFA as antigenically relevant, maps closely onto HA1 sites with the highest ratio of non-synonymous to synonymous mutations, with most lying in epitopes (the 25 amino acid positions with the highest significance are listed in S3 Table), providing an improved view of the physio-chemical constraints at play in influenza antigenic evolution [20]. Furthermore, we uncover a significant amount of antigenically relevant interactions in H3 hemagglutinin, as suggested previously [21, 22]. We show that the importance of many amino acid positions depends on the time frame of the training set (Figure 3), with some locations being significant throughout. The predicted RFA significance for each amino acid is quite closely related to the total number of non-synonymous mutations observed at those positions, reinforcing the suggestion that positive selection is operating on a relatively small subset of HA1 amino acids – S3 Figure. This apparent association of accumulation of non-synonymous mutations with increased phenotypic plasticity deserves further consideration. Examination of the genetic evolutionary trajectories of epitopic and non-epitopic sites in HA1 reveals that if HA1 amino acids are mutating exclusively under neutrality assumptions (using a Kimura model with 2:1 transition to transversion ratio and equal mutation probability for each position), the simulated viruses generally increase their differences in polarity and hydrophobicity when compared to their ancestral HK69 strain in a liner fashion. Framing those neutral expectations with the observed trends, we see that while HA1 as a whole has evolved near neutrally (S4A Figure), epitopes have undergone strong positive selection, optimizing their polarity and hydrophobicity differences relative to the ancestral HK68 virus (S4B Figure).

**Figure 3.**
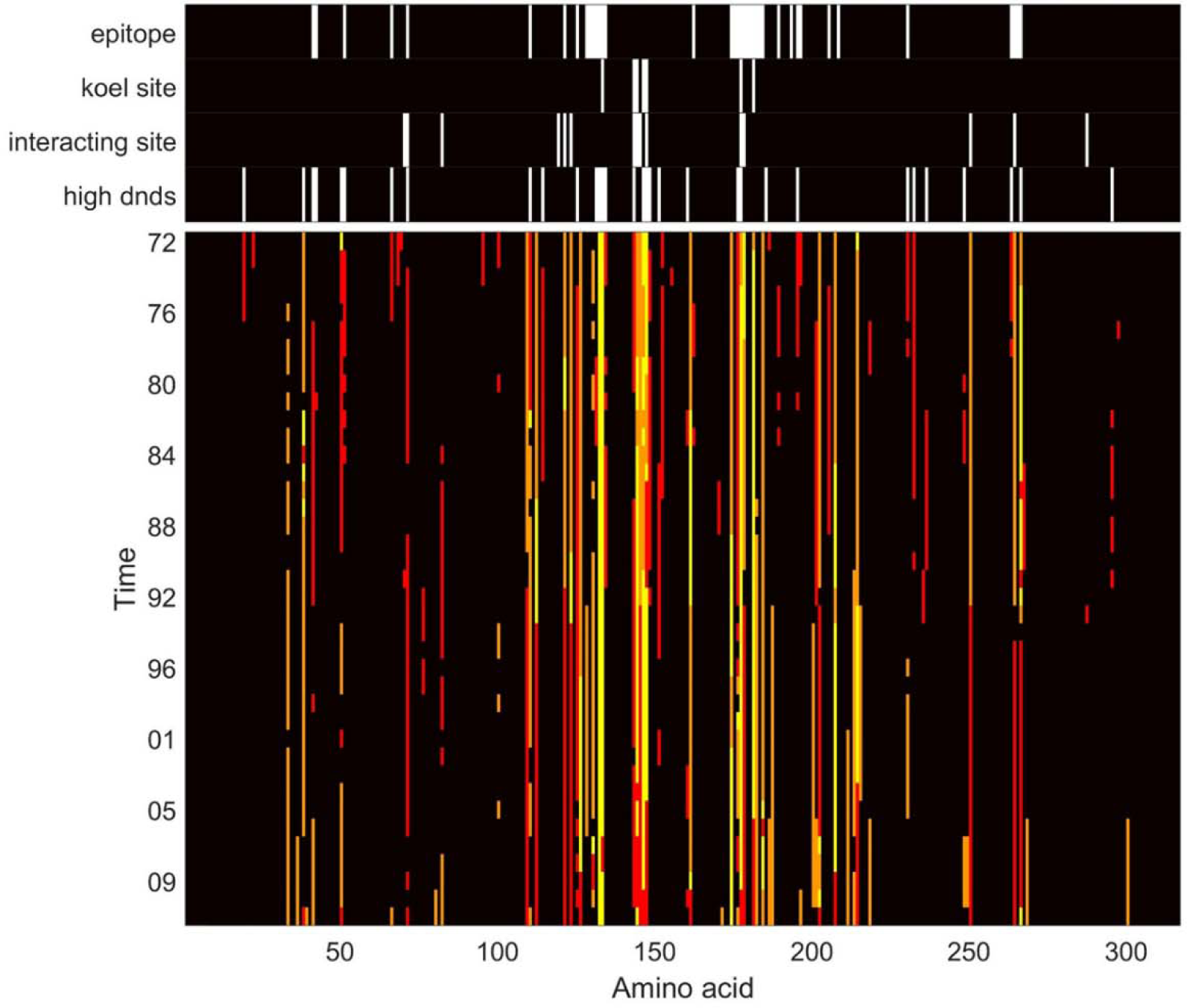
Mapping the positions with significance in explaining the observed antigenic distances between antigen/sera pairs of sequences. RFAs were performed using training data encompassing different time spans. If a particular HA1 position wa significant in an RFA trained on data collected from a specific time point onwards, that position is colored orange. If that position is significant in the RFA trained on data from 1968 up to that time point, the position is colored red. If the position is significant in both of those time frames, it is shown in yellow. The significant positions in the RFA map closely to positions with a high ratio of non-synonymous to synonymous mutations, with most occurring on known HA1 epitopes (highlighted in white above).

It seems then that there are a few key amino acids in HA1 displaying a larger than expected phenotypic plasticity, evident in the range of observed polarity and hydrophobicity differences as well (S5 Figure), which seem to be under positive selection. A detailed analysis of how mutations are distributed across HA1 reveals a surprisingly stable pattern, with approximately 50% of all mutations across any antigen/serum pair falling on this subset of highly phenotypically relevant sites identified by the RFA – S6 Figure. Other epitopes are under-represented, whilst about a third of mutations lies on non-epitopic polymorphic sites.

### Exploring evolutionary trajectories and the limits of antigenic evolution predictability

Having demonstrated that it is possible to predict the antigenic characteristics of new H3N2 strains several years into the future with a good but not perfect level of accuracy (Figures 2 and S2), we now consider whether it is possible to predict which strains will arise in the future. The linearity of the antigenic map suggests the existence of a strong immune-mediated selection compelling novel influenza viruses to maximize their antigenic distance to previously circulating viruses [3, 10]. Above, we show how this is manifested as an uneven distribution of non-synonymous distributions across HA1, resulting in accumulation of differences in physio-chemical properties at key sites. Furthermore, a look at how genetic and antigenic evolutionary trajectories match, indicates that both antigenic and genetic differences are much smaller within members of the same cluster than between members of adjacent clusters (S7 Figure). This suggests there is a limited number of genetic evolutionary trajectories which translate into cluster jumps, but does not help resolve whether the observed antigenic evolutionary trends are a result of:

– structural constraints limiting the physio-chemical plasticity of new viruses (“plasticity”).
– immunological selection pressure, forcing new viruses to accrue physio-chemical differences relative to circulating ones (“antigenic filtering”).
– seeding effects, with apparent “canalization” of antigenic and genetic evolutionary trajectories resulting from the expansion of a single viral lineage in a susceptible population (“seeding”).
– All the above.

Here, we consider how such constraints shape the joint genetic/antigenic evolution of H3 and how they translate into the possible evolutionary trajectories the virus might follow. We try to investigate these effects independently to address their significance in recovering the observed antigenic maps, specifically trying to reproduce cluster jumps. To do so we use a genetic simulator algorithm coupled with a GP map and compare the predicted antigenic distances between simulated viruses and both the ancestral strain and sampled viruses belonging to the following antigenic cluster. In our genetic simulator we explore 3 molecular evolutionary models, expressing quite different mutation distribution profiles: a) 7-site model, in which mutations are restricted to the sites identified as cluster determinants by Koel and colleagues [23]; b) RFA 25-site model, where mutations are restricted to the 25 most significant amino acid positions as predicted by the random forest trained on the full dataset; c) Generalized model where mutations are distributed according to probabilities directly derived from S6 Figure.

Whilst the latter model is in agreement with the ‘canalized’ antigenic evolution [10] and the antigenic recycling [8] hypotheses, since the majority of mutations is directed at a small subset of positions, it also allows for new, previously antigenically irrelevant, alleles to become significant [19] and epistatic interactions between antigenically significant sites and other loci [24].

We first compare the accuracy of the GP map obtained when including all sites in the training set with that of the antigenic maps resulting from antigenic distances predicted by RFAs trained exclusively on the two limited sets of sites listed above. The 25-site RFA model performed comparably to the generalized model in reproducing the training set and in terms of predictive accuracy unlike the 7-site model that, while broadly capturing cluster transitions, was substantially poorer at reproducing the precise size and shape of antigenic clusters (S8 Figure). Thus, while 7-site may be sufficient to explain much about cluster transitions, our analysis suggests that more sites contribute to the fine detail of antigenic variation within clusters, and consequently, finer level antigenic novelty leading to antigenic cluster transitions.

To ascertain the validity of our joint GP map and molecular sequence generating tool, we simulated thousands of viruses from a single SY97 cluster ancestor using the generalized evolutionary model and attempt to replicate the antigenic location of the FU02 cluster. Given perfect information as to what positions mutate in the future and using GP map trained on the complete dataset we can almost exactly retrieve the observed antigenic map (S9 Figure). Whilst still intercepting the antigenic space occupied by the empirical future cluster, blinded predictions reflect a degree of uncertainty as to the antigenic trajectory undertaken by flu viruses, with a range of antigenically distinct viruses equally likely to emerge. Indeed, less than 1% of the simulated viruses were within 2 units of antigenic distance of the empirically realized FU02 cluster. Looking deeper into the genetic relationship between simulated sequences and historic ones, we can see that only a minority of the antigenically plausible sequences are genetically very similar to the observed FU02 antigenic cluster (S10 Figure).

It then seems that “antigenic filtering” alone cannot reconcile genetic and antigenic H3 evolution, so we next examine what roles “plasticity” and “seeding” might have. To do so, we compare how the expected mean antigenic distance of simulated *n*-point mutants to their ancestral strain, and between themselves fits with those observed in the empirical data. We introduce “plasticity”, “antigenic filtering”, and “seeding” effects sequentially, thus always limiting the physio-chemical plasticity of each HA1 site and stacking the other effects in succession. We start with a strain in the SY97 cluster as the ancestor (see S6 Figure for another starting point) and train the RFA only on strains from SY97 and earlier clusters. Figure 4A illustrates how the predicted antigenic distance between simulated viruses with a specific number of mutations to the ancestral strain compares with the predicted antigenic distance between pairs of equally evolved simulated viruses (with same number of mutation to the ancestral). The mean antigenic distance between simulated viruses is consistently larger than their respective antigenic distances to the common ancestor if no “antigenic filtering” is imposed, i.e. in the absence of immune pressure. When antigenic selection is introduced (Figure 4B), the simulated viruses somewhat converge antigenically, being, on average, more similar to each other than to viruses in the ancestral cluster. This effect is more a consequence of the forced increase in the mean antigenic distance between simulated viruses and their ancestor resulting from the immune selection filter (S11 Figure) than a true antigenic convergence, since there isn’t a significant decrease in the pairwise antigenic differences between simulated viruses. This would suggest that selection for antigenic novelty could result in any one of many antigenically distinct viral lineages. Whilst these results are clear for both the generalized and 25-site models, they are much less so for the 7-site model (S11 Figure) due its antigenic sensitivity to changes in a very small number of sites. Although mutations can occur in up to 25 amino acids in the later model, only changes in 7 of those actually result in an antigenic effect which then tends to be quite strong (these sites have all been associated with cluster jumps), making general extrapolations on the effect of immune selection difficult with this model.

**Figure 4.**
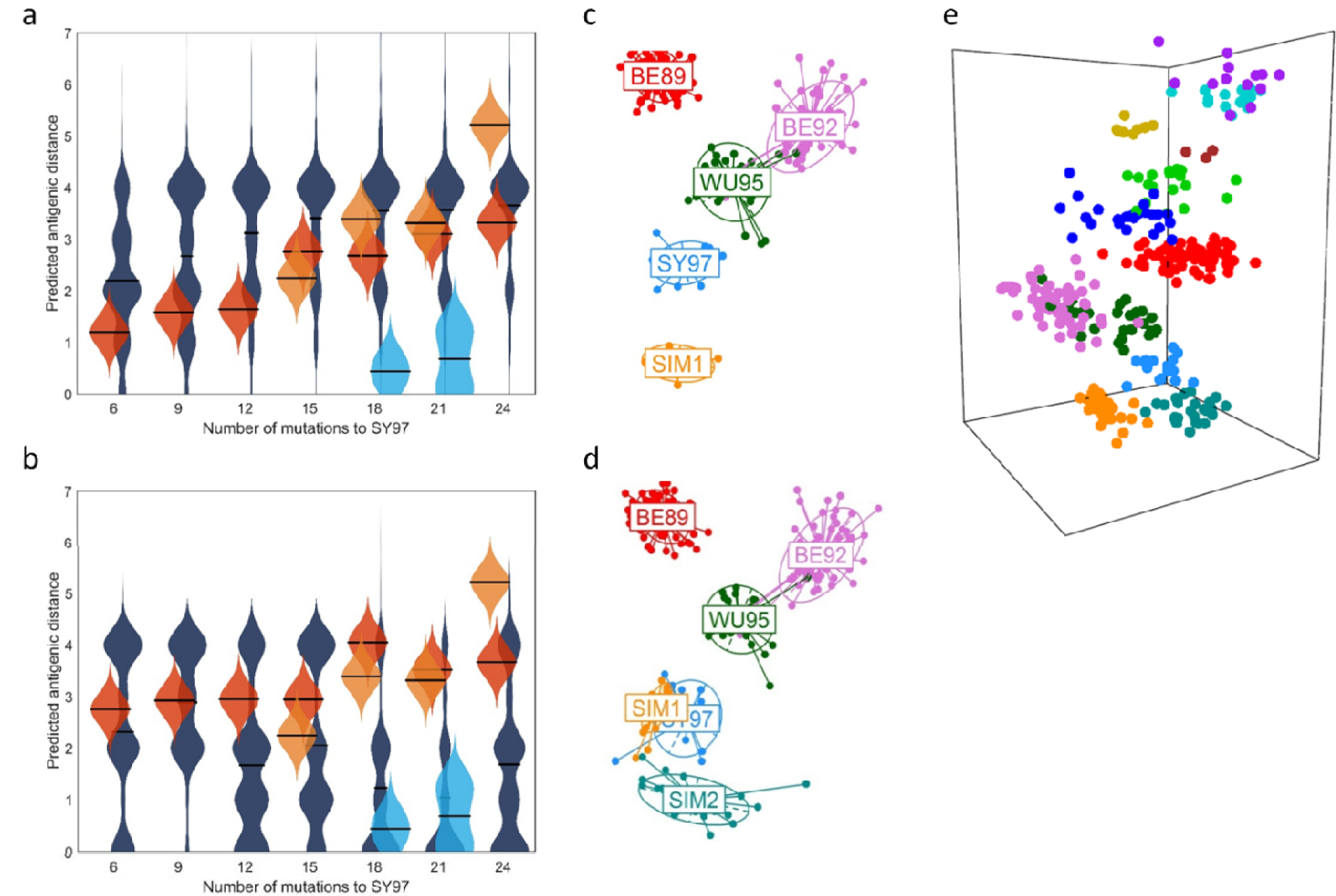
Unfiltered (A) and filtered step-wise (B) antigenic evolution from a given ancestral strain. The violin plots illustrate the kernel density estimates for two predicted antigenic distances (horizontal lines represent the mean) on sets of simulated viruses with X amino acid mutation relative to an ancestral strain (a member of the SY97 cluster). The dark blue violin plots represent the distribution of antigenic distances (with the respective means as horizontal lines) between simulated viruses with X amino acid mutations relative to a virus in SY97. The red violin plots give the predicted antigenic distance across simulated viruses. Light blue violin plots give the empirical antigenic distance between viruses in SY97 and FU02, whilst the orange plots refer to the measured antigenic distances within the FU02 cluster. (C) Antigenic map of one simulated antigenic cluster arising from SY97 (see S1 Text for description of simulation methods). (D, E) Antigenic maps including two simulated clusters both 3-5 antigenic distance units away from their SY97 ancestor, on different spatial embeddings. The 2-Dimensional embedding is unable to properly resolve the antigenic distances between the simulated clusters in contrast with the 3-Dimensional representations (E).

When we simulate a gradual evolutionary process akin to what has been proposed previously [25–27], with multiple related lineages co-circulating globally with occasional expansion of one of those lineages in temperate locations following migration, here termed ‘seeding’, we observed much stronger antigenic convergence. Quite strikingly, the mean predicted antigenic distance between simulated viruses only matches the observed mean difference among FU02 cluster samples when this stepwise process was implemented (Figures 4B, S8).

### Predicting cluster jumps and implications for vaccine updates

If ‘seeding’ is indeed a critical process, we need to better characterize which viruses can serve as seeds to a cluster jump. We found that any one of the intermediate simulated viruses resulting from the second mutational step could seed an antigenic cluster which would look nearly identical to the FU02 cluster on an antigenic map (Figure 4C). This is not an indication of the similarity of this simulated cluster with the FU02 cluster but rather a reflection of how the geometry of antigenic maps is mainly determined by large antigenic distances between observed viruses and their antisera. The similar antigenic map location of the observed FU02 cluster and several simulated clusters (we only show a representative one) is solely a result of those clusters sharing the same distance to their ancestor. If two new lineages were to emerge simultaneously, the resulting map would show substantial antigenic separation between them (Figure 4D). We find that the accuracy of the 2-dimensional map when reproducing pairwise distances between antigen and sera substantially worsens when mapping multiple simulated antigenically divergent clusters of viruses, with a 3-dimensional representation being preferred (Figure 4E). The adequacy of a 2-dimensional representation thus results from the single lineage seen in H3N2 HA evolution, rather than necessarily being the fundamental underlying dimensionality of the antigenic shape space which influenza evolution explores [28] – S12 Figure.

We show that we can reproduce the observed antigenic patterns of relatedness among simulated viruses and between simulated and sampled viruses, as well as identify viruses that are antigenically novel and likely seeds of new clusters. Next, we push the boundaries of evolutionary predictability by trying to prospectively replicate observed cluster jumps. To do so, we take a late sample (collected the year prior to the establishment of a new cluster) of a given antigenic cluster and simulate viruses according to the generalized model, including all the evolutionary constraints listed above (‘plasticity’, ‘filtering’, and ‘seeding’). The GP map uses a RFA which is trained on data only up to the year of collection of the strain used as a seed, but here we also calculate distances to sampled viruses in the future cluster, thus comparing how predicted clusters fit with the realized antigenic trajectory.

Strikingly, all simulated cluster jumps reveal the same patterns, with 3 quite distinct antigenic trajectories being theoretically possible, out of which only one resembles the observed cluster (Figure 5). We can identify specific mutations generating pitchforks in the antigenic trajectory followed by simulated viruses – see S1 text for more details. From the set of 6-14 positions highlighted as significant to explain antigenic lineage segregation, there is consistently at least one that has previously been identified (19) as a cluster jump determining mutation – S13 Figure.

**Figure 5.**
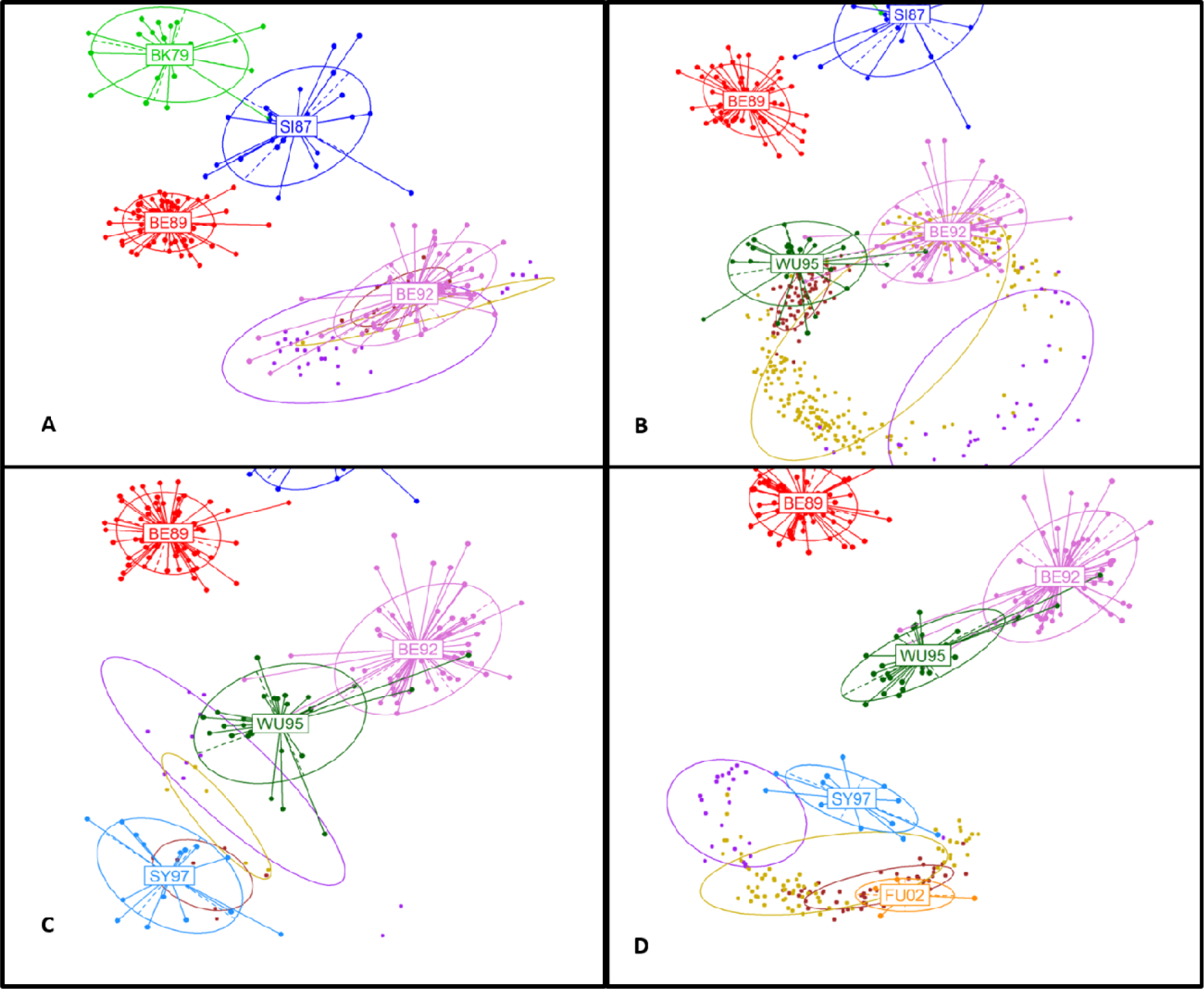
Evolutionary trajectories in antigenic space. The panels illustrate 4 cluster jump predictions. From a given cluster an RFA is trained on data up to a specific dat when the chosen ancestral strain was collected, and 100,000 viruses are genetically evolved according to the generalized model. The simulated viruses are classified into distinct groups according to their respective predicted antigenic distances to the observed future cluster and all preceding clusters. The simulated viral groups (or antigenic lineages) are colored purple, yellow and brown in descending order of antigeni similarity to the observed future cluster, respectively.

## Discussion

Here, we have shown how machine-learning based feature selection methods can be used to generate an explicit genotype to phenotype (GP) map that can reproduce the observed features of A/H3N2 influenza antigenic evolution and offers insights into its predictability. Our results indicate that the antigenic distance of newly isolated viruses from their ancestors can be accurately predicted from their HA1 sequence with the methods presented here. This suggests that such GP maps could usefully supplement experimental antigenic characterization of newly sampled viruses in routine virological surveillance for vaccine strain selection.

However, the predictability of phenotype from genotype [11,13,23,29] does not imply influenza evolution is fundamentally predictable with a narrow set of possible antigenic trajectories the virus can undertake at a point in time. In contrast, our analysis suggests that influenza‘s observed evolutionary pathway is but one of a limited set of antigenic trajectories, with mutations in antigenically relevant sites (particularly HA1 epitopes, but not exclusively) generating a breadth of antigenically novel viruses later subject to epidemiological selection pressures which limit overall viral diversity [6,10,27]. We conclude that selection for immune escape does not tightly constrain influenza’s antigenic evolutionary trajectory but rather that epidemiological selection (with marked winter epidemics and summer bottlenecks) coupled with strong seeding effects (largely determined by the existing intense global mobility network) determine the observed antigenic trajectory. For all the counterfactual antigenic trajectories to be observed, a much more intense and constant sampling effort would be required.

Clearly there are limitations to the use of GP maps, in that they can only be trained on what has been observed. While we partially mitigated this limitation by training our RFA model on changes in physiochemical properties of HA amino-acids rather than the amino-acid sequences directly, mechanistic models predicting the conformation and antigenicity of HA from its sequence would clearly be preferable. However, these remain a long-term goal. Our work indicates that machine-learning derived GP maps, coupled with judicious animal studies [23], are now able to give substantial scientific insight into the phenotypic evolution of influenza viruses as well of being of practical benefit in enhancing virological surveillance.

## Materials and Methods

### Genotype-to-phenotype (GP) map

We trained a Random Forest Algorithm (RFA) on a dataset of pairwise differences in physiochemical properties between the amino acid sequences of antigen/serum pairs and the respective antigenic distances measured by the HI assay. The RFA is an ensemble classifier consisting of multiple low correlation decision trees, which aggregate into a low bias and low variance “forest”. Each tree in a random forest is trained on a random subset of the data, and each tree branch contains randomly chosen variables from all available variables (in this case, a subset of positions in the amino acid sequences for each branch). Final classification of each sample results from aggregating the votes of all trees in the forest, with the importance measure for each variable being the loss of classification accuracy caused by the random permutation of attribute values for that variable. Overall, RFA displays excellent performance in classification tasks [30, 31] and provides direct measures of both variable importance and classification error. We used the RandomForest R package to implement the RFA. For more information see S1 Text.

### Antigenic mapping

Antigenic mapping [3, 28], by offering a simple intuitive representation of large volumes of HI assay data, is a useful tool for unveiling the antigenic evolution of a pathogen. It is fundamentally a multidimensional scaling (MDS), and more specifically, a metric unfolding problem (due to the sparseness of the data). Assuming the empirical HI data is directly related to Euclidian distances in an antigenic space, we can view the problem of map creation as that of the minimization of an error function, *E*:

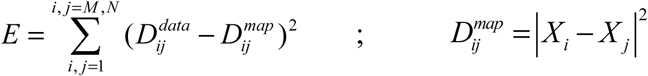

where 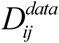 correspond to the measured antigenic distances, and 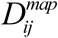 to the Euclidean distances computed between the location, *X_i_*, of the *i^th^* of *M* antigens in shape space, and *X _j_*, the location of the *j^th^* of *N* antisera.

To create an antigenic map we choose an embedding dimension, *d*, and then fit the *d*(*N*+*M*-1) parameters specifying the locations of the *N* antigens and *M* sera (fixing one location at the origin). The map thus created not only provides a visual interpretation of measured antigenic distances, but also implicitly predicts unmeasured distances antigen/serum pairs. See S1 Text for more information on our implementation of a maximum-likelihood based multidimensional scaling algorithm for antigenic mapping.

### HA1 genetic evolutionary model

Molecular evolution simulations are rooted on a mutational algorithm that generates non-synonymous mutations in specific codons with a 2:1 transition to transversion ratio, following Kimura’s nucleotide evolution model [32]. In practice, a given number of mutations is assigned to a particular subset of amino acid positions and for each mutation up to 2 nucleotides from the respective codon are chosen to undergo mutation (keeping with a 2:1 the transition to transversion ratio). The procedure is repeated if necessary until one gets a non-synonymous substitution. This algorithm was implemented in Matlab® R2016a. We can thus generate hundreds of thousands of simulated viruses from any sampled H3 virus. We apply this DNA mutation model to 3 different genetic evolutionary frameworks expressing quite different mutation distribution profiles ranging from a model where mutations are restricted to only 7 amino acids [23]; to one where mutations are allocated to the 25 most significant amino acid positions as predicted by the RFA; and a generalized model where mutations are assigned to different amino acids according to a 4:1:2 ratio (RFA significant sites/other epitopes/other polymorphic sites).

### HA1 antigenic evolutionary model

Throughout this paper we try to ascertain the determinants of observed H3 antigenic evolutionary trends and consider the roles of physio-chemical plasticity (“plasticity”), immunological selection pressure (“antigenic filtering”), epidemiological seeding effects (“seeding”).

The imposed physio-chemical plasticity is bounded by what is observed in the data, with mutations in simulated viruses having to comply to the observed minima and maxima in each amino acid position for the considered physio-chemical properties – S5 Figure.

Since the mean antigenic distance between viruses in two adjacent antigenic clusters is 4.46 (±1.3) on average), simulations complying to the “antigenic filtering” effect impose boundaries on simulated viruses’ predicted antigenic distances to past clusters, according to two simple rules: 1) the simulated viruses must have predicted antigenic distance values within the 3-5 range relative to all viruses within the reference cluster; 2) predicted distances to viruses in all other past clusters must be greater than 6. We thus discard any simulated viruses predicted to have an antigenic distance to viruses of the reference antigenic cluster greater than 5 or lower than 3, and simultaneously a predicted antigenic distance lower than 6 to members of any other past cluster. We implemented the “seeding” effect as a 3-step mutational process (4-6 mutations in each step) where at the end of the 1^st^ and 2^nd^ steps one simulated virus is selected as a seed for the next simulation step.

## Acknowledgments

We thank the EU FP7 EMPERIE project, the NIGMS MIDAS programme and the MRC for research funding.

## Author Contributions

RA and NMF conceived the study. RA implemented the genotype to phenotype map and developed the molecular simulation algorithm. RA and NMF wrote the paper.

---

**S1 Figure.**
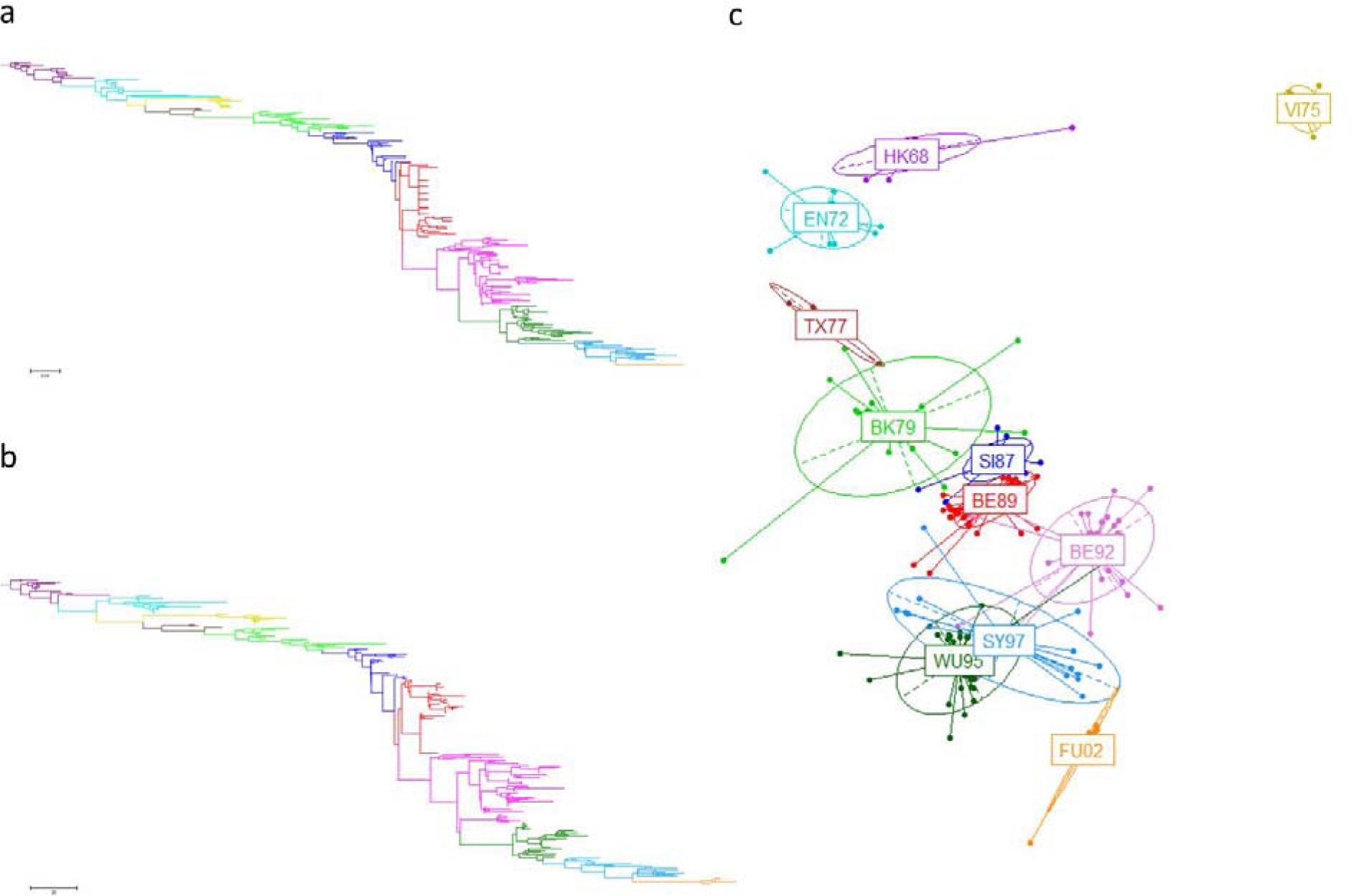
Correspondence between genetic distances and sum in polarity and hydrophobicity across the sequences. (A) ML phylogenetic tree of all antigen sequences. (B) Dendrogram built from a distance matrix derived from the sum of differences in polarity and hydrophobicity in the antigen sequences. (C) MDS using the same matrix as in (B).

**S2 Figure.**
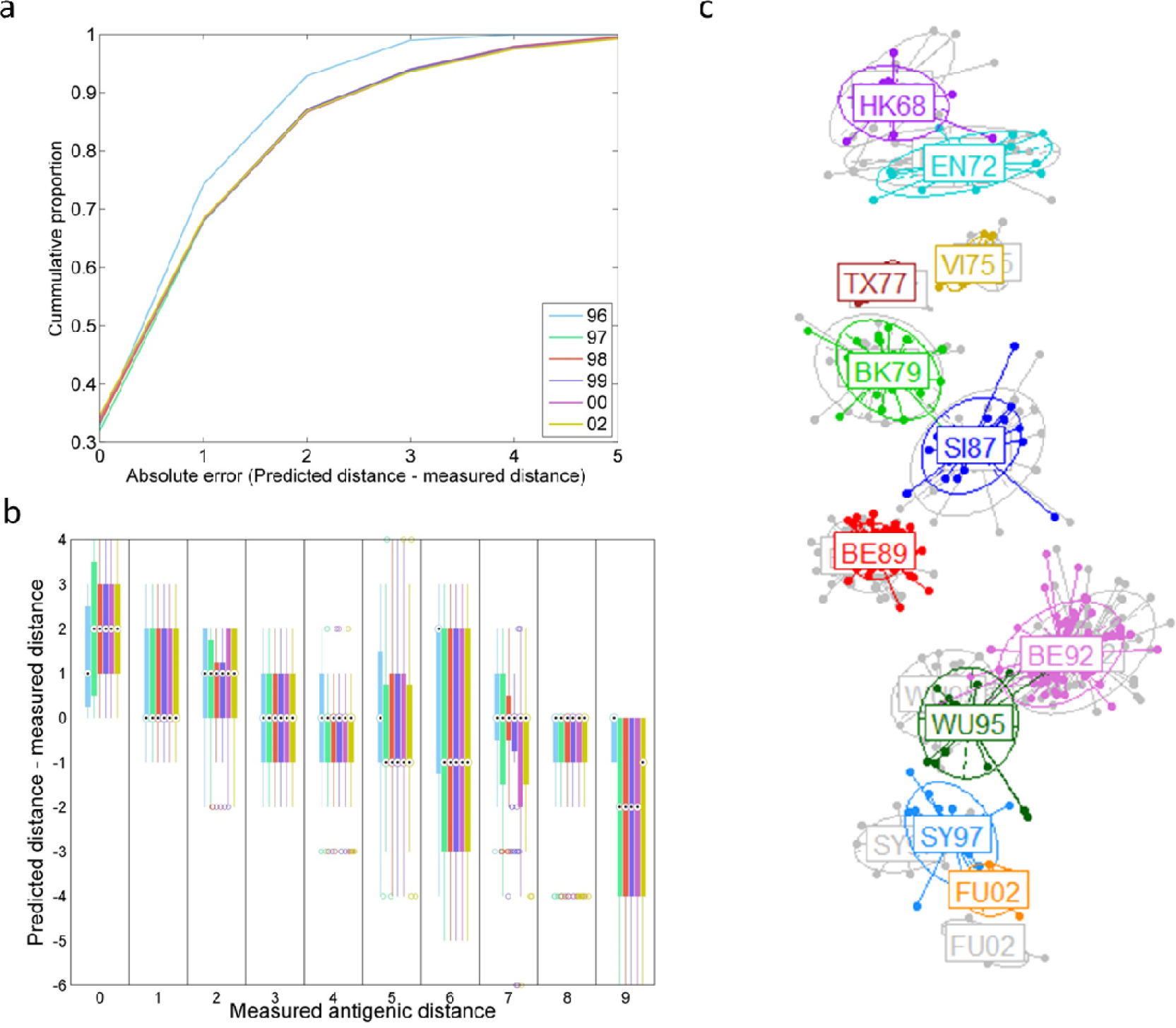
Prospective predictive power of the RFA algorithm with predictions made from a training set with data collected up to 1995. (A) Cumulative curves representing the distribution of the prediction errors (on a log-2 scale) for the different prediction time windows. (B) Box plots of the prediction error for specific values of measured antigenic distance. For each box, the central mark represents the median, the edges mark the 25^th^ and 75^th^ percentiles, and the whiskers extend to the most extreme data points not considered outliers. Here the whiskers extend to +/–2.7*variance (corresponding to 99.3 coverage assuming the data are normally distributed). Each colour represents an incremental prediction time window as in (A). (C) Predicted antigenic map with predictions in colour overlaid on the map generated with full hindsight (in grey).

**S3 Figure.**
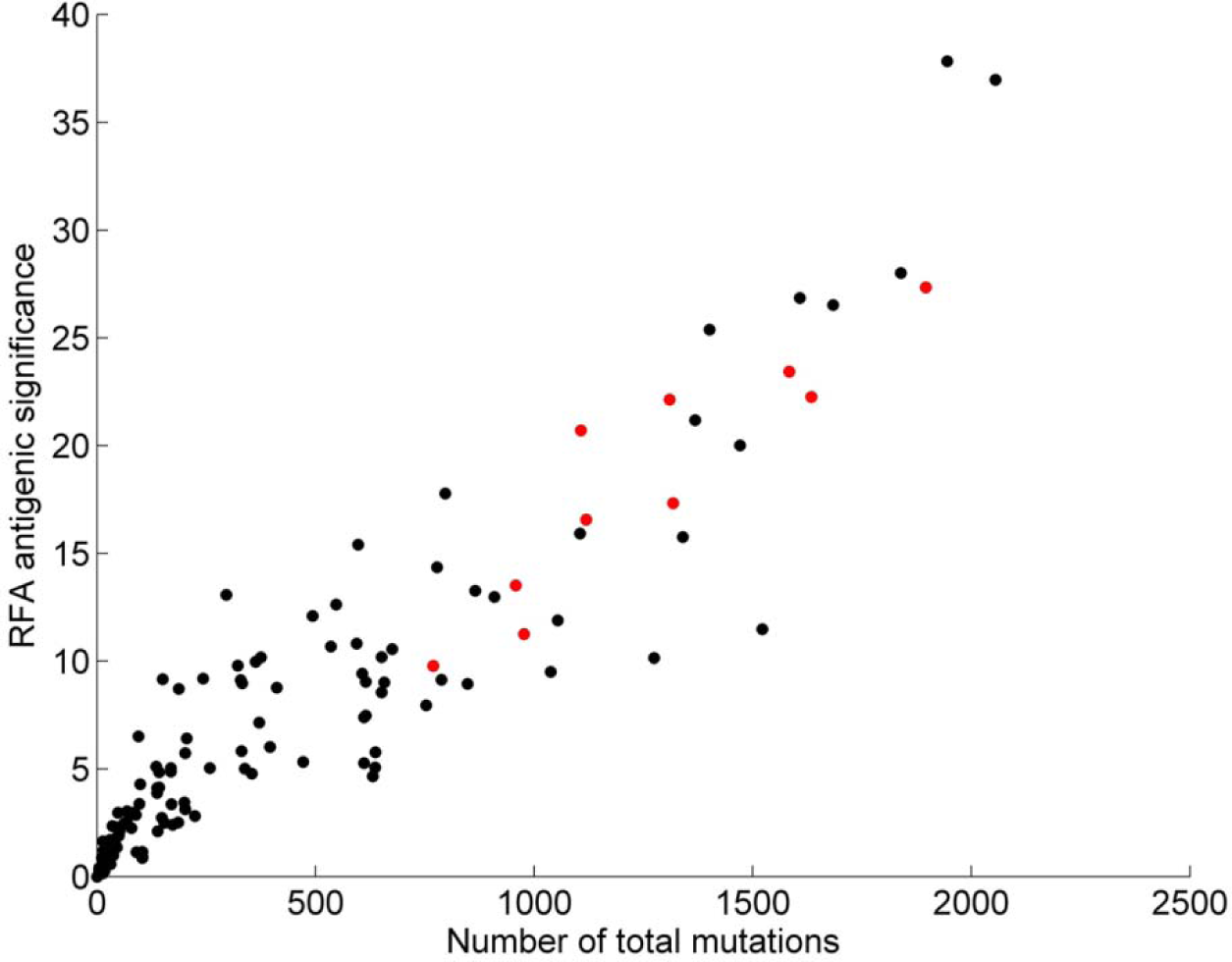
Correlation between the total number of mutations recorded at each position and the respective RFA (trained on the full dataset) significance. Amino acid positions with high interaction indices are highlighted in red.

**S4 Figure.**
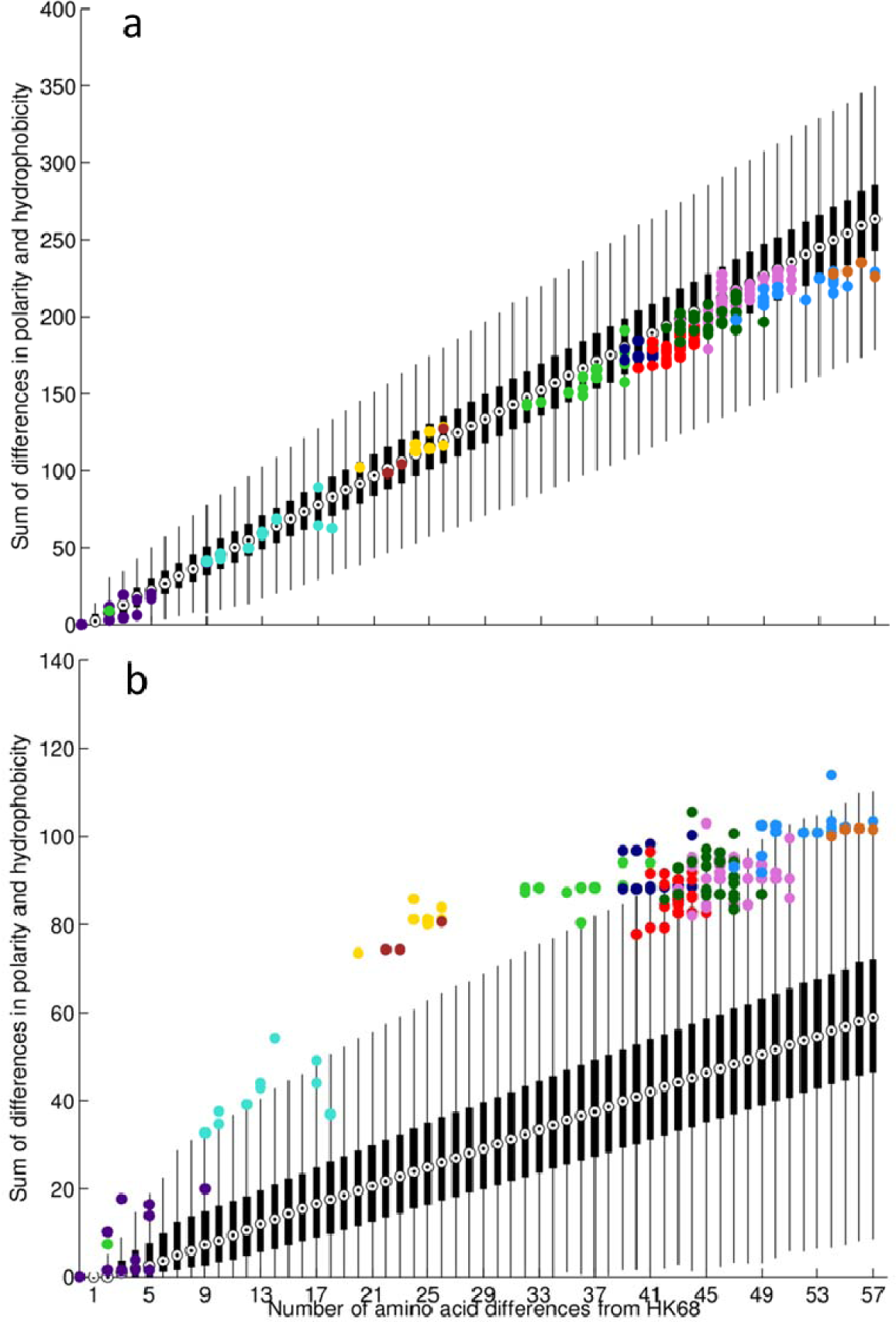
Evolution in HA1. Each box plot in black show the variation in the sum of differences in amino-acid polarity and hydrophobicity between 100,000 viruses and their ancestral HK68 virus (given a specific number of mutations). The black circle in each box identifies the median, the edges mark the 25th and 75th percentiles, and the whiskers extend to the most extreme data points not considered outliers. (A) Shows differences across the full HA1 sequence, and (B) across epitope positions only. Isolated viruses are color-coded according to their assigned antigenic cluster.

**S5 Figure.**
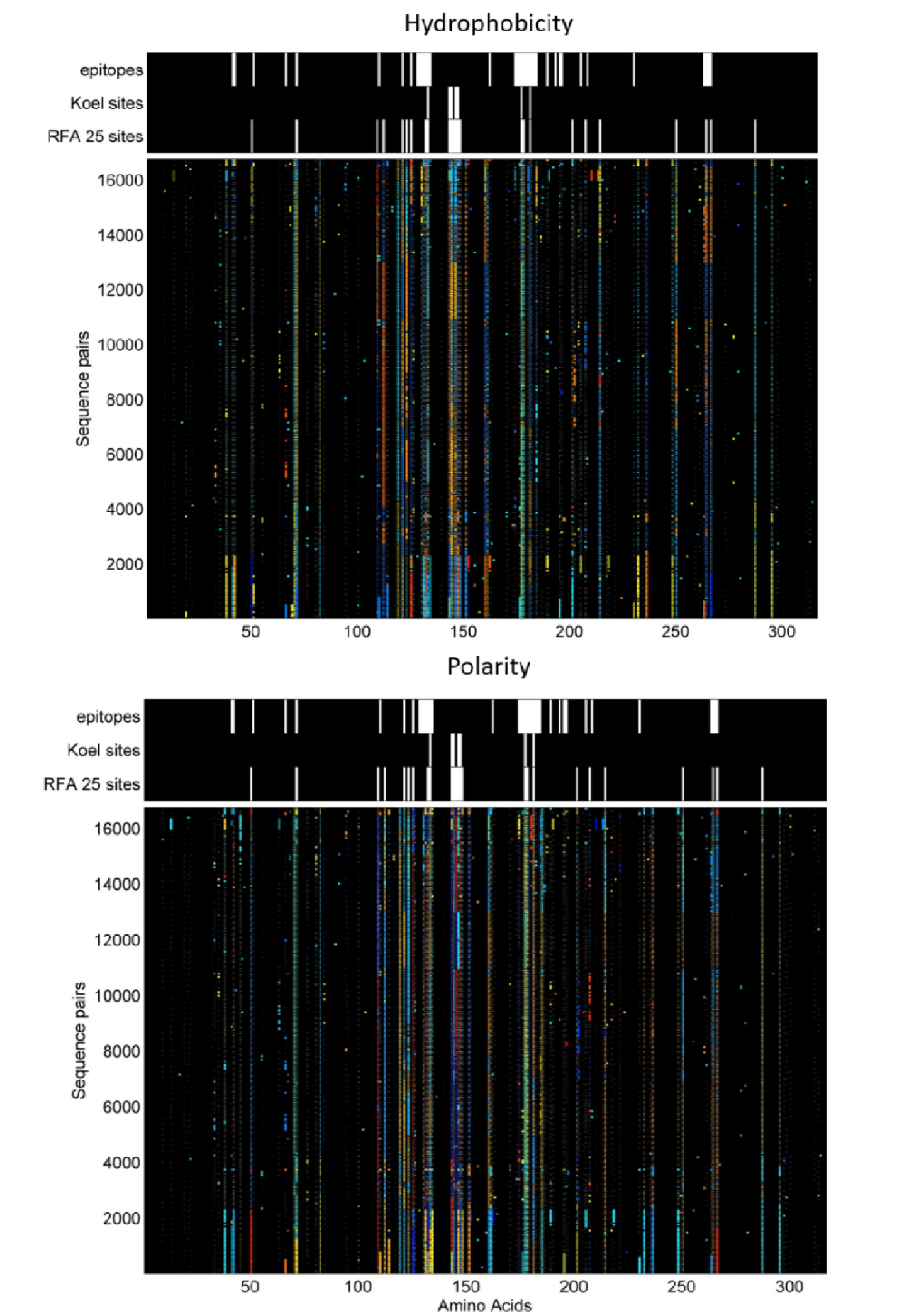
HA1 physiochemical plasticity. Displays the observed difference in hydrophobicity (top) and polarity (bottom) for all antigen/serum pairs (rows) in each amino acid position (columns). Black indicates no difference, whereas blue and red represent the most extreme values (negative and positive, respectively).

**S6 Figure.**
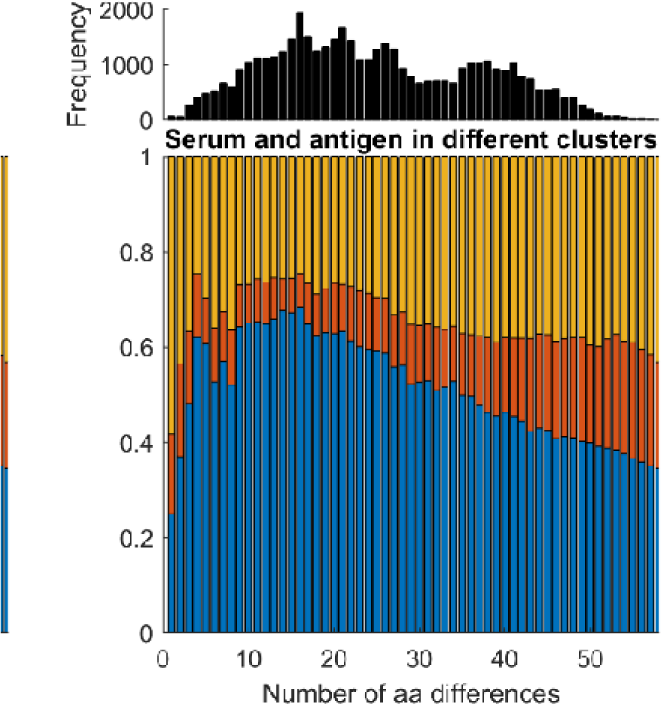
HA1 mutational profile. Shows the distribution of mutations across three subsets of amino acids. Blue illustrates mutations that fall on the subset of RFA 25 most significant sites, in red we represent the known epitopes which are not included in the blue subset, and in orange the remaining polymorphic sites (not included in any of the other subsets).

**S7 Figure.**
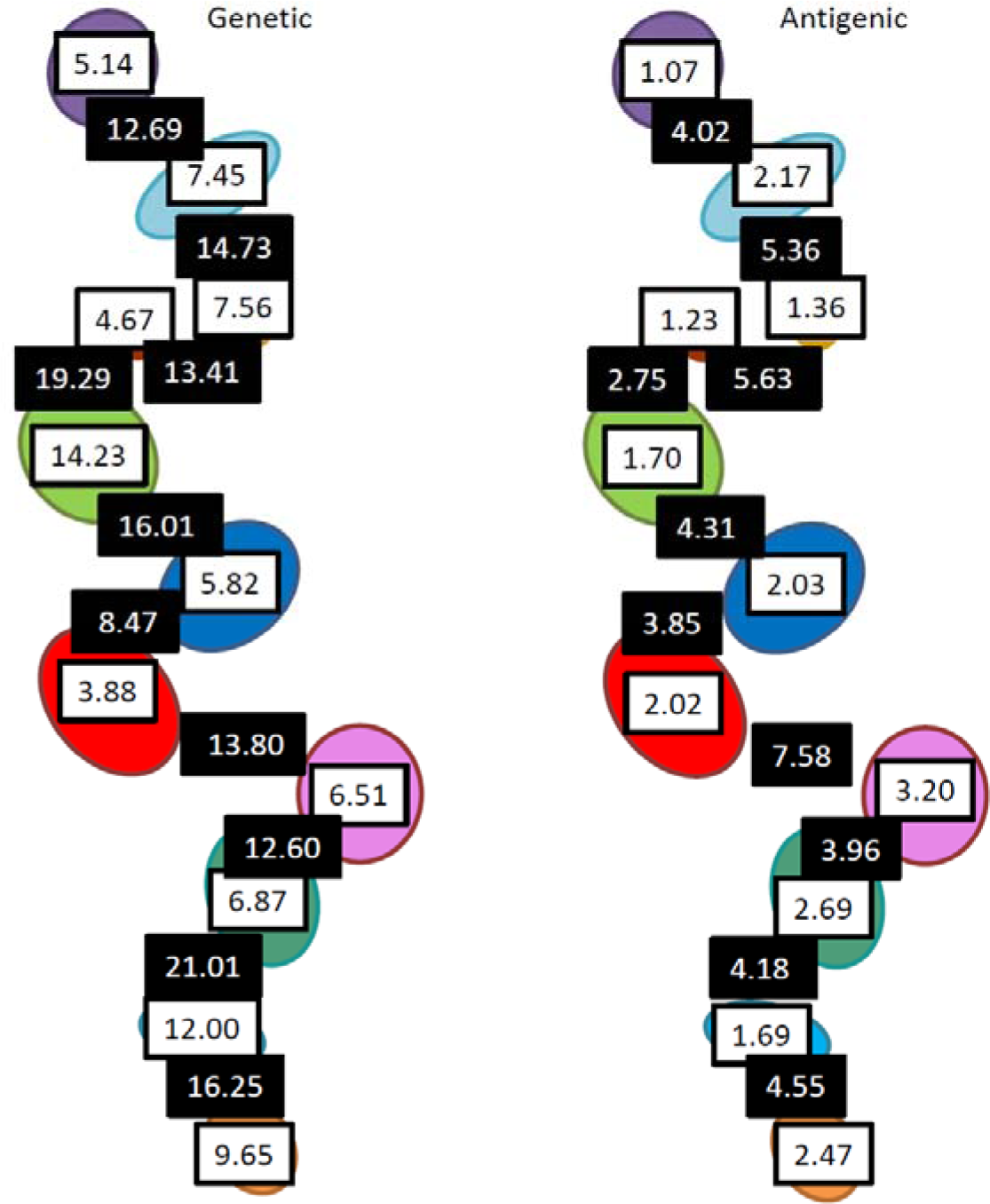
H3 Antigenic and Genetic Evolutionary patterns. This sketch illustrates mean differences between members of the same cluster (white background boxes) and between viruses belonging to contiguous clusters (black background). The mean antigenic differences shown here are calculated from the HI titers measured in the considered antigen/sera pairs. The mean genetic differences are merely the mean number of amino acid differences between sequences of the same cluster or across contiguous clusters.

**S8 Figure.**
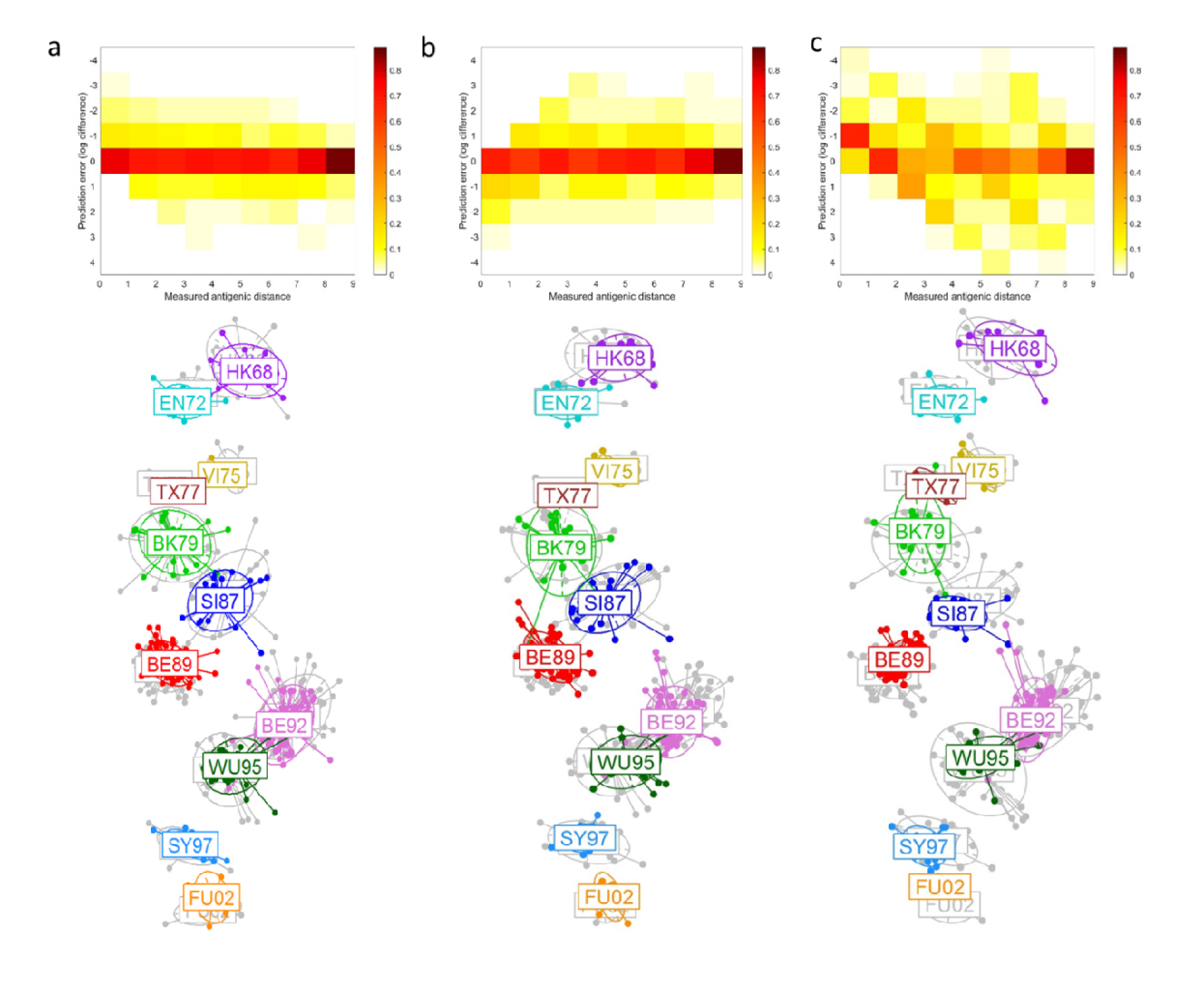
Evolutionary model accuracy. Classification accuracy of the RFA looking at hydrophobicity and polarity differences in each of the considered evolutionary models: the generalized (A); the RFA 25 most significant sites (B) or the 7 sites model proposed by Koel *et al* (C). Superimposed 2-dimensional antigenic maps generated from the predicted antigenic distances (in colour) - when training the RFA on the respective subsets of amino acids - and from the HI titre data (in grey). The top panels give the prediction errors for each of the observed antigenic distance bins.

**S9 Figure.**
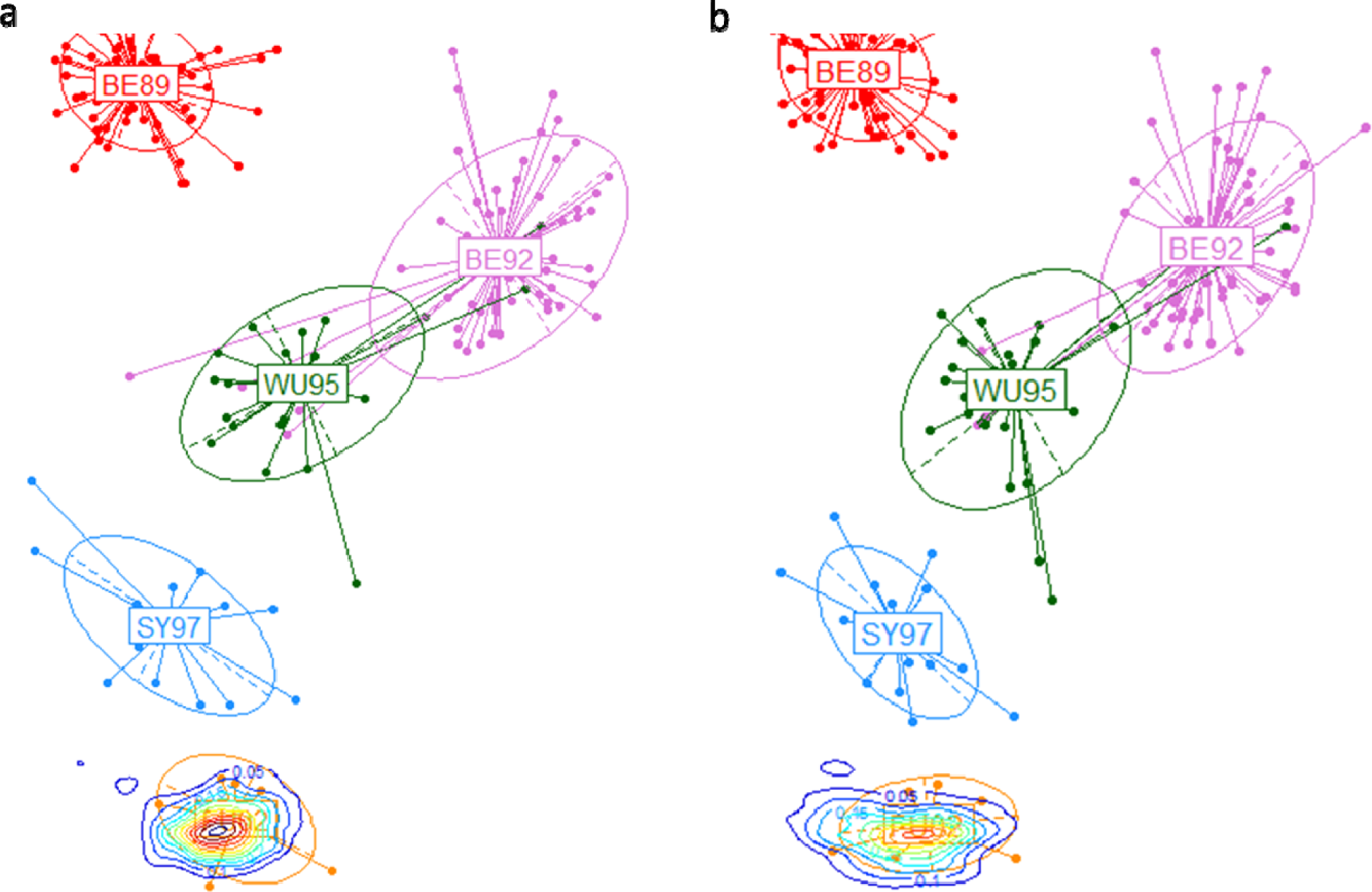
Antigenic maps of influenza’s predicted antigenic trajectories. The contour line represent the density probability that simulated viruses will fall on the respective delineated antigenic space. The simulated viruses were generated from mutations in: (A) significant positions to explain antigenic differences between clusters BE92 to FU02; (B) the polymorphic sites across the SY97and FU02 clusters. Each simulated virus has a total of 12-18 amino acid mutations relative to its ancestor.

**S10 Figure.**
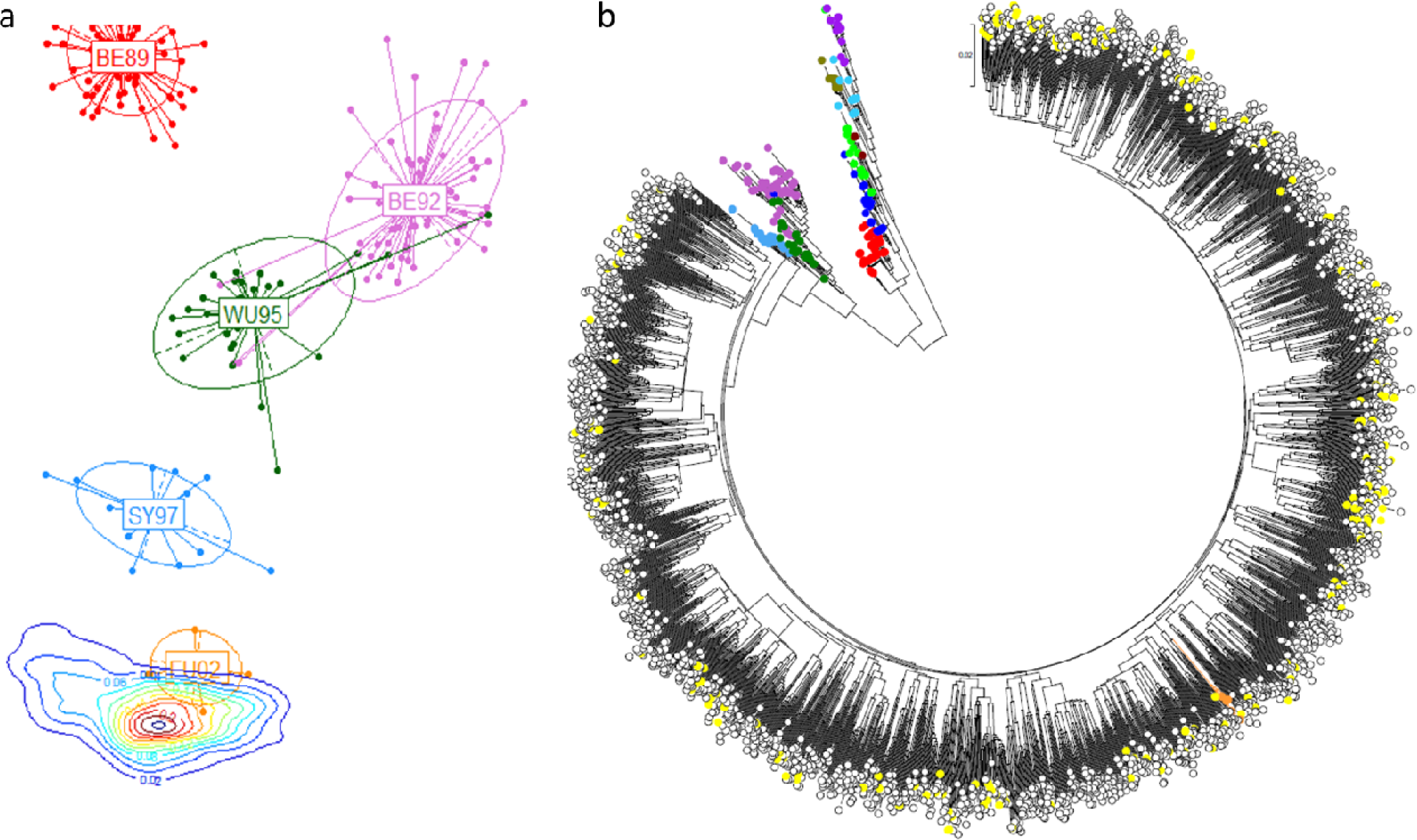
Antigenic trajectory unpredictability. Antigenic map of influenza’s predicted antigenic trajectory using the generalized model (A). We implemented a mutational process similar to that in S9 Figure and impose an antigenic filter that only accepts simulated viruses with predicted antigenic distances to the ancestral viral cluster in [3, 5] and of over 6 to all preceding clusters. The contour lines represent the density probability that simulated viruses will fall on the respective delineated antigenic space. A maximum likelihood neighbor joining tree shows the genetic relationship between simulated viruses that did not meet the antigenic filter requirements (white), those that did (yellow), and the empirical viruses (colored according to their antigenic cluster as in all other plots). Note the empirical FU02 cluster in orange amidst the simulated viruses.

**S11 Figure.**
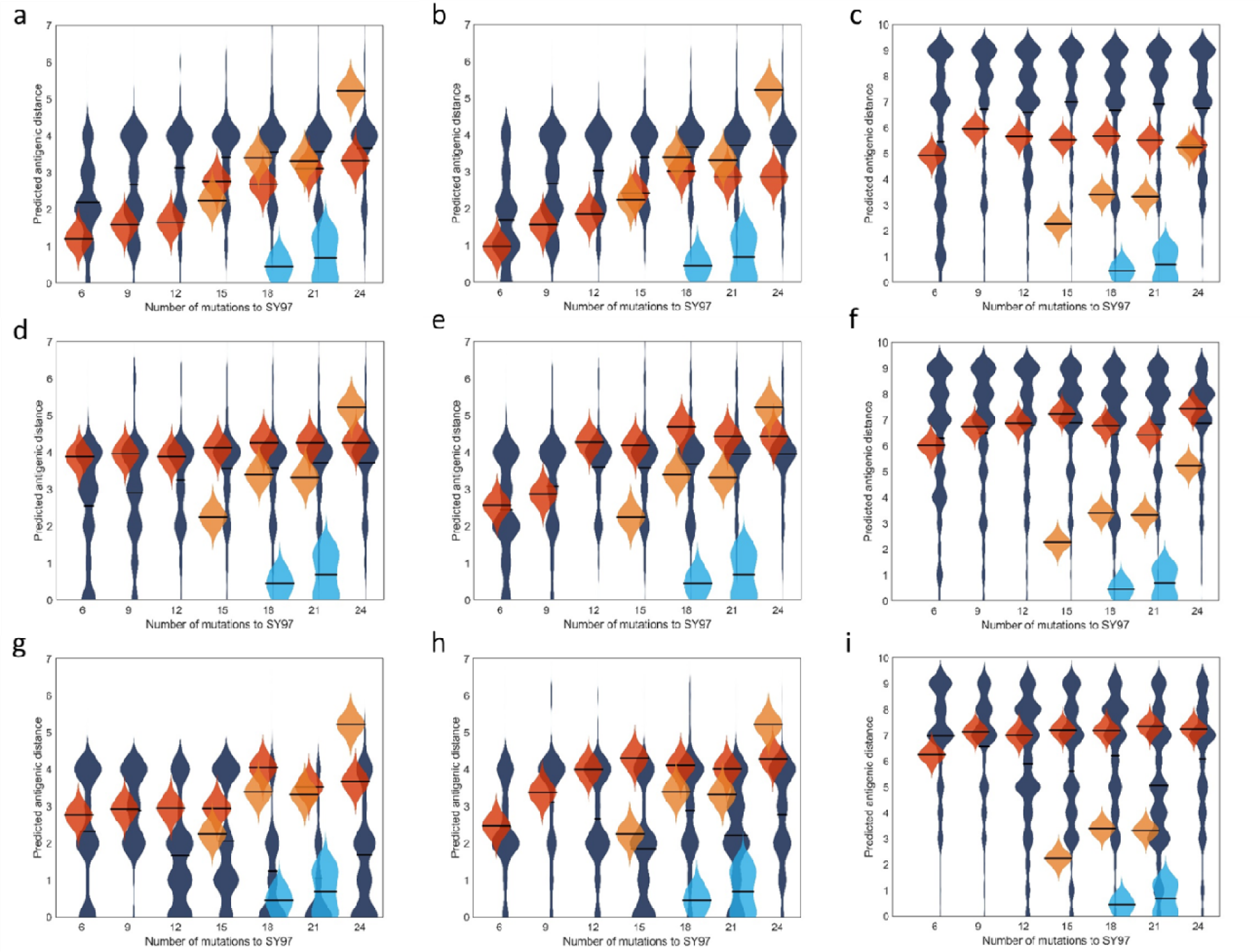
Antigenic trajectories and underlying evolutionary processes. Unfiltered (top row); filtered (middle row); and filtered step-wise (bottom row) antigenic evolution from a specific ancestral strain under the 3 evolutionary models. The results for the generalized, 25-site, and 7-site models are depicted left to right respectively. The dark blue violin plots represent the distribution of antigenic distances (with the respective means as horizontal lines) between simulated viruses with X amino acid mutations relative to a virus in SY97. The red violin plots give the predicted antigenic distances across simulated viruses. Light blue violin plots give the empirical antigenic distances between viruses in SY97 and FU02, whilst the orange plots refer to the measured antigenic distances within the FU02 cluster.

**S12 Figure.**
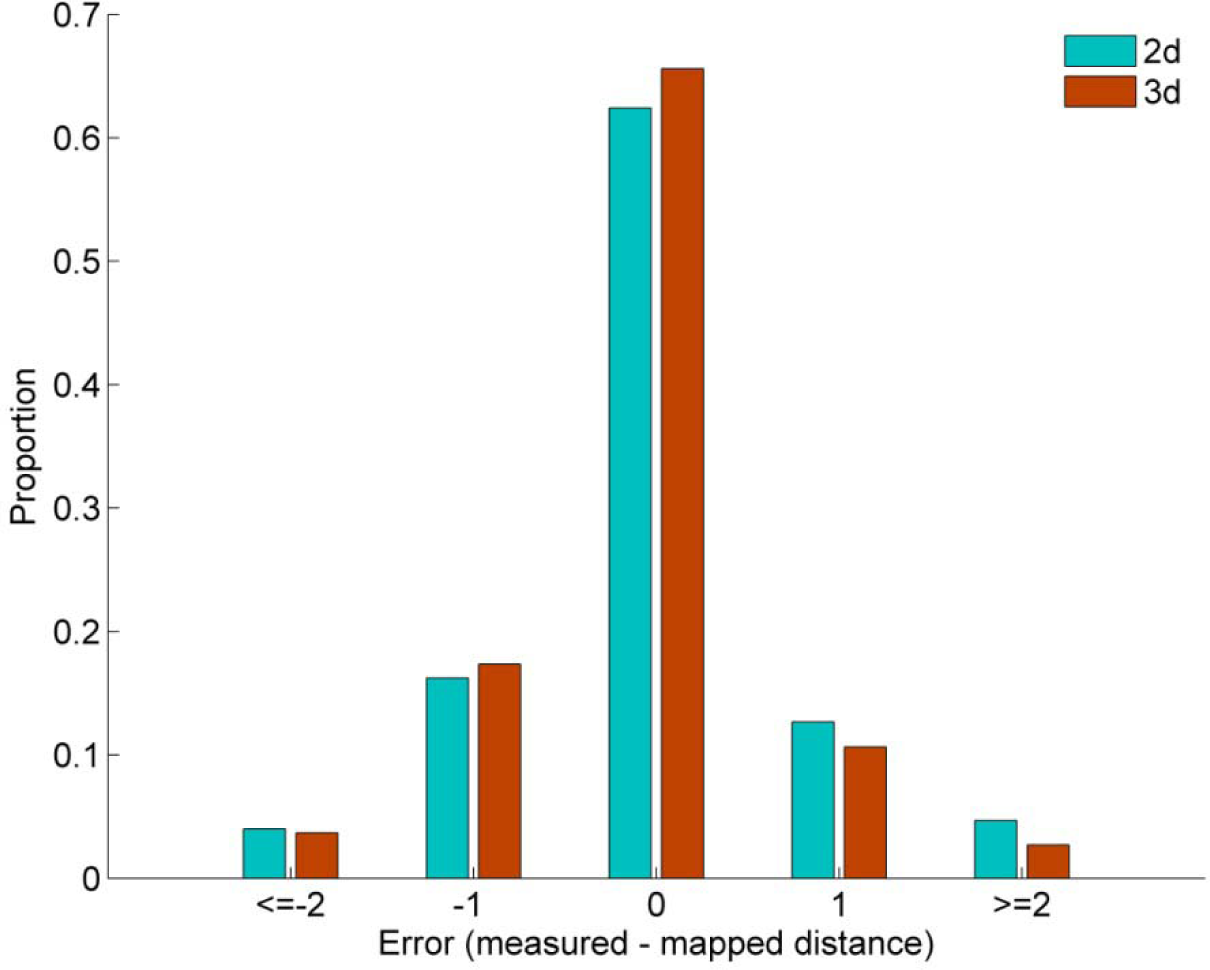
Antigenic map precision on different embeddings. Comparison of the MDS error for the 2D and 3D representations of an antigenic map with two simulated lineages.

**S13 Figure.**
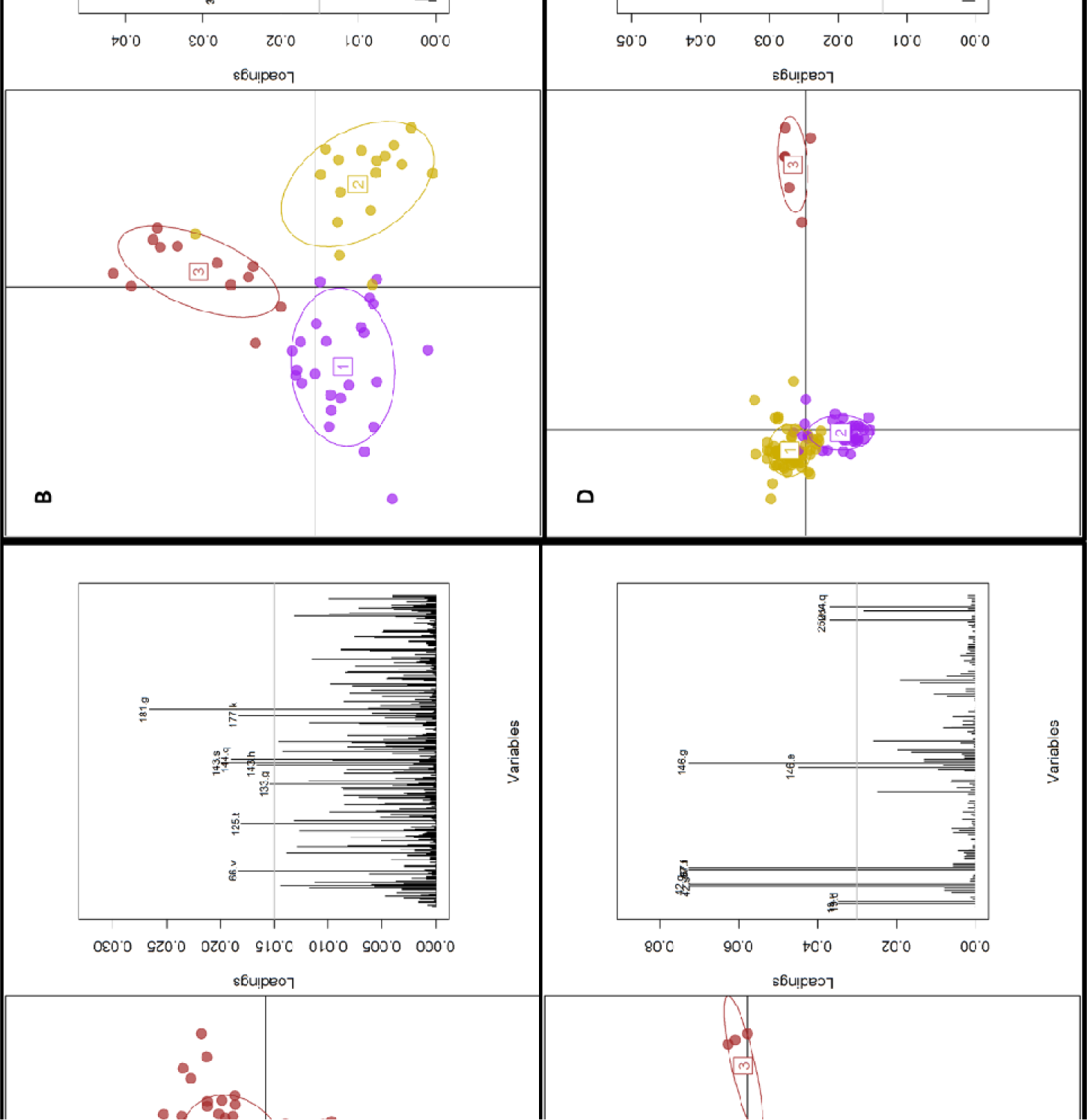
Genetic determinants of simulated viral lineages. Each panel relates to the associated cluster jump in Figure 5 of the main text. The left plot of each panel shows the DAPC resulting discrimination of the separate groups given the genetic information. The contribution of each allele for the segregation of the viruses into groups is shown on the plots to the right. Th dashed line serves only as an arbitrary threshold above which the individual allele labels are shown.

**S14 Figure.**
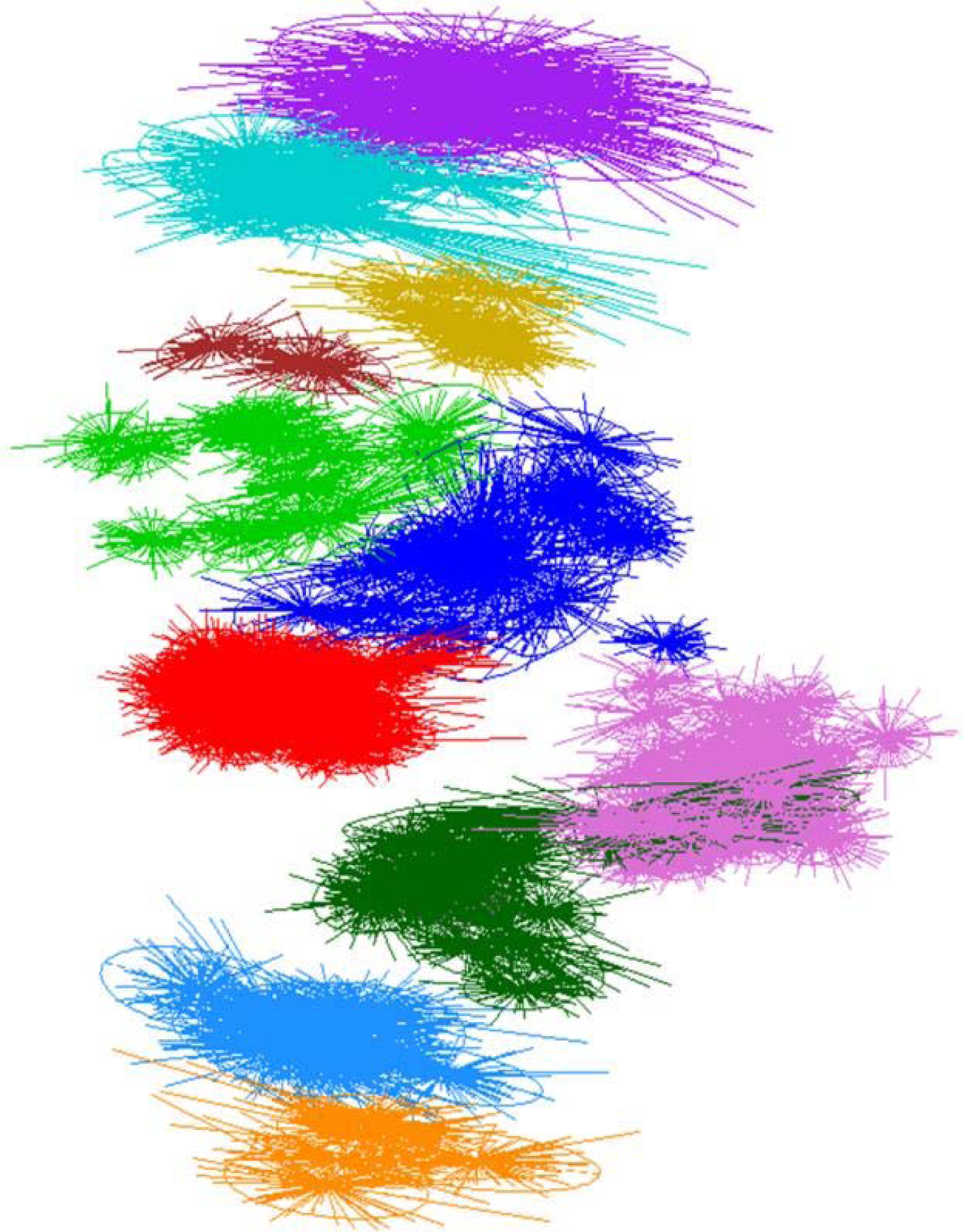
Uncertainty of the antigenic mapping algorithm of historic H3 influenza. Each ellipsis represents the dispersal in the final proposed coordinates for each antigen (over 100 bootstrap runs of the mapping algorithm). Colors match the cluster assignments in.

**S15 Figure.**
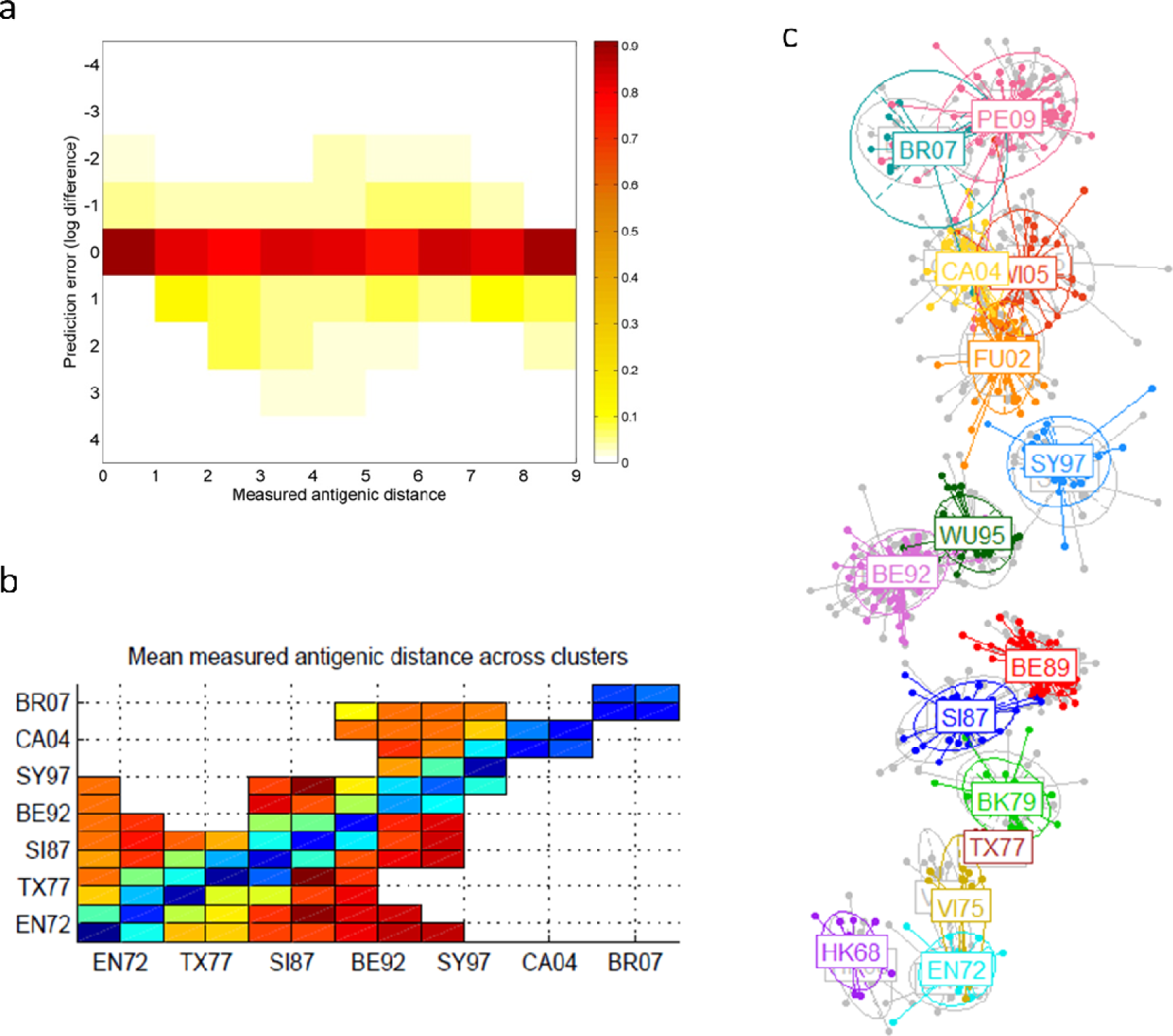
Classification accuracy of the RFA using hydrophobicity and polarity as antigenic proxy variables for an extended dataset. (A) Surface plot showing the error in prediction of antigenic distance (distance being measured on a log-2 scale) for every antigen-serum pair. Antigen-serum pairs are grouped by measured antigenic distance on the horizontal axis. The colour of each rectangular pixel shows the proportion of pairs with at a particular measured antigenic distance with a particular prediction error. (B) Mean measure antigenic distance between antigen/serum pairs composed of elements of different antigenic clusters. The sparseness of this matrix of measurements increases significantly for more recent data. (C) Superimposed 2-dimensional antigenic maps generated from the RFA predicted antigenic distances (in colour) and from the HI titre data (in grey).

