## Supplemental Text and Tables for "Predictability of the antigenic evolution of human influenza A H3 viruses"

Supporting Information

#### *The data*

We used the hemagglutination inhibition (HI) assay data published in Smith *et al*.[1], filtering out those antigens and sera for which the complete HA1 sequences are not available in the GenBank database. Antigens and sera composing the HI data panel used throughout are shown in Extended Data Table 1. Following past work [2], we defined the antigenic distance between antigen *i* and serum *j* to be:

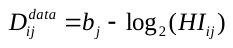

where *HI_ij_* is the measured HI assay result for antigen *i* and serum *j* and *b_j_* is the *log*_2_ of the maximum HI measurement for antiserum *j*.

The HI assay outputs a viral titer which indicates the number of times the sera needed to be diluted for hemagglutination of the antigen and red blood cells to occur [3]. This suggests that there is an electrostatic interaction between antigen and antibodies that is disrupted once the serum is too diluted. Given this premise, we set out to determine whether we could use physiochemical properties as proxy for the antigenic distances observed in the HI assays. To do so, we implemented a machine learning algorithm that was informed on how differences in 54 distinct physiochemical properties relate to empirical HI titre measurements – more discretized description of the method can be found in the Genotype to Phenotype Map section below. Out of those 54 properties, a subset of 7 (*α* helix, *β* sheet, *β* turn, bulkiness, coil, polarity and hydrophobicity) emerged as the most relevant in explaining the observed phenotype, with polarity and hydrophobicity being by far the ones with the most significance.

S1 Figure illustrates how well differences in polarity and hydrophobicity represent the HI distances between the H3 viruses; we see these give a surprisingly accurate representation of influenza’s antigenic clustering. To generate this figure we calculated the sum of the absolute differences in polarity (*pol*) and hydrophobicity (*hydro*) – values for amino acid hydrophobicity and polarity as specified by the Aboderin [4] and Grantham [5] scales, respectively – for each antigen/serum pair (*ij*), for each amino acid in HA1 (*p*):

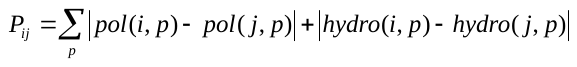

These differences constitute a dissimilarity matrix which can directly be used to build a dendrogram that can be compared with a phylogenetic tree of amino acid differences.

Given these results, we assumed differences in polarity and hydrophobicity could represent antigenically significant physiochemical differences across two viruses and then use these differences (rather than the underlying amino acid sequences) to train our GP map. Use of physiochemical properties have been previously used for functional analysis [6] and antigenic characterization [7].

Although a more up to date collection of HI data is now publicly available [8], the matrix of antigen/serum HI measurements for viruses collected after 2003 is much sparser than that the earlier dataset. In particular, distances are only recorded for viruses in contiguous (in time) antigenic clusters, resulting in very poor anchoring of the antigenic maps and much larger uncertainty in absolute antigenic positions than that observed in the dataset analyzed here – S15 Figure. Given the generality of the antigenic evolutionary trends explored in this paper we have therefore chosen not to include that most recent HI data.

#### *Genotype to Phenotype Map*

A number of rather different technical approaches have been implemented to develop a genotype to phenotype map for influenza A virus antigenicity. Some approaches used regression methods to establish the relevance of certain pieces of genetic information in determining antigenic distance [9–12]. Others have adopted cluster analysis to identify phenotypic clusters arising from specific genetic comparison metrics [13,14]. Phylogenetic analysis has been used to build antigenic trees and evaluate the antigenic impact of amino acid changes [15]. Lastly, protein folding algorithms have been used to inform a neutral network model [16].

Here, we opt for a machine learning approach that correlates differences in physiochemical properties across all amino acids of a pair of viral sequences with a measure of antigenic distance between them. This method has the advantage of accounting for possible interactions between loci which is critical since in traditional genome wide association studies (GWAS), single nucleotide polymorphisms (SNPs) typically only explain a fraction of the phenotypic variance [17,18].

Sophisticated statistical techniques attempting to discriminate phenotypically relevant variables (disease phenotype, gene expression patterns) have become more prominent in biomedical research over the past decade. These feature selection methods attempt to highlight the subset of relevant features that need to be included to achieve accurate classification [19,20]. One such algorithm is the random forest algorithm (RFA) which has been recently adapted to the analysis of microarray data [21,22] and host specificity of emerging viruses [23]. It offers excellent performance in classification tasks, and provides direct measures of variable importance and classification error, whilst identifying a subset of nucleotide or amino acid positions that give the most discriminating information regarding the phenotype of interest. As an embedded feature selection technique, RFA has several advantages over other types of algorithm. Embedded methods incorporate variable selection as part of the training process making them more effective because they make integral use of the training data (whereas other methods require a split into training and validation sets) and incorporate feature selection as a part of the model training process [24].

RFA is in essence an ensemble classifier consisting of multiple low correlation decision trees, which aggregate into a low bias and low variance “forest”. Each tree in a random forest is trained on a random subset of the data, and each tree split contains randomly chosen variables from all available variables (in this case, a subset of positions in the amino acid sequences for each split). Final classification of each sample results from aggregating the votes of all trees in the forest, with the importance measure for each variable being the loss of classification accuracy caused by the random permutation of attribute values for that variable. The prediction error of the RFA is calculated by the ‘.632+’ bootstrap method which relies on the relationship between the resubstitution error and the misclassification error on samples not used to train the algorithm (out-of-bag samples). The resubstitution error essentially measures the proportion of observations in the original dataset that are misclassified by the decision trees within the random forest. Variable importance is also estimated from the samples which are left out of the training set at each split of the tree, making the random forest algorithm very robust to over-fitting. In fact Random Forests have been demonstrated not to over-fit as the number of trees grows to infinity, instead producing a limiting value of the prediction error [22].

We train an RFA on a dataset composed of pairwise differences in physiochemical properties (polarity and hydrophobicity) between the amino acid sequences of antigen/serum pairs and the respective antigenic distances measured by the HI assay. In practical terms, the algorithm takes as inputs *n* sequence pairs of size *k*p* (*p* being the number of amino acids in each HA sequences, and *k* the number of evaluated physiochemical properties), and a class variable *h* defining the antigenic distance between the respective antigen/serum pair. The algorithm then tries to find a subset
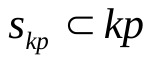
of amino acid positions whose contribution to the explanation of the class variable is significantly greater than that of random permutations of the values of variables
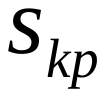
 in the out-of-bag samples. In practice, we associate each HI measurement with the differences in polarity and hydrophobicity observed at each amino acid of the respective sequences. The RFA randomly selects a few of those measurements (for each tree) to serve as a training set and outputs measurements for the significance of each property’s change for each amino acid, that can then inform a regression model to accurately predict the antigenic distance for the samples outside the training set. The overall significance of a specific amino acid position *i* is given by
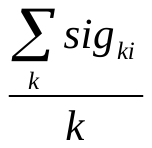
.

#### *Antigenic Maps*

Antigenic cartography serves as a useful tool for unveiling the patterns of antigenic evolution of a pathogen. Antigenic maps, particularly two dimensional ones, offer an intuitive way of representing the information contained in extremely large matrices of antigenic distance measures (in HI titer values), such that HI values in the original distance matrix are monotonically related to the Euclidian distance between the corresponding points in the newly created shape space. Assuming the empirical HI data is directly related to Euclidian distances in space, we can view the multidimensional scaling problem in terms of the optimization of an error function, *E*:

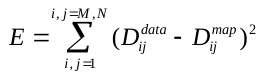
 ;
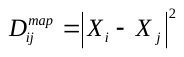

where
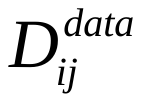
correspond to the empirical distances, and
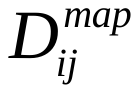
to the Euclidean distances computed given the coordinate vectors, *X*, of the points in two dimensional space, with
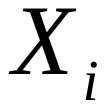
 representing the position of the *i^th^* of *M* antigens in shape space, and
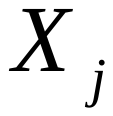
the location of the *j^th^* element of *N* antisera. We implemented a Markov Chain Monte Carlo (MCMC) approach to estimate the locations of antigens and sera on 2 dimensions such that *E* (taken as our log likelihood) is minimized. To do so, we use the Metropolis-Hastings algorithm, perturbing the location of all antigen and sera on both axis (*X*) by sampling from a Gaussian distribution with mean *X* and evaluate the changes in likelihood. For each iteration *t* we:

- Propose candidate locations *X^*^* by sampling from the Gaussian distribution *G*(*X^*^|X_t-1_*).
- Calculate the *acceptance ratio* 
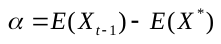
 used to inform acceptance or rejection of the new proposal.
- If
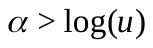
, *u* being a uniformly distributed random number between 0 and 1, *X^*^* is accepted. Otherwise it is rejected and
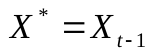
.

We ran the mapping algorithm 10 times using the final proposed coordinates as initial conditions for the subsequent run and used the final proposals of the 10^th^ run as initial conditions for all the maps depicted throughout. The algorithm runs for 100 iterations, with *i*j* proposals in each iteration. For each antigen/serum pair we propose candidate locations for that antigenic and that serum and evaluate the consequent acceptance ratio.

The stability of the mapping algorithm solutions can be evaluated by looking at the displacement of the final coordinate proposals across several runs. S14 Figure depicts the spatial displacement of each antigen location across 100 of the mapping algorithm runs which use the same initial conditions but different random number generation seeds. Each ellipsis represents the variance in the final proposed coordinates of each antigen and is colored according to the cluster to which that antigen was assigned in [1].

The antigenic maps thus created provide not only a visual interpretation of the empirical antigenic distances, but also a complete depiction of influenza’s antigenic evolution patterns by inherently providing distances for unmeasured antigen/serum pairs.

#### *Making antigenic distance predictions*

With our GP map we can take any new viral sequence and predict the antigenic distance of that virus from any sequence used to train the map. To assess the applicability of our RFA to newly collected viruses, we compare the observed distances in measured antigenic distances between viruses collected after a specific time point to antigenic distances predicted after training the RFA with data only up to that reference date. S2A and S2B Figures illustrate the resulting out-of-sample prediction accuracy for a GP map trained with data collected up to 1995 with predictions being made on antigen/serum pairs where at least one of the elements was collected after 1995. The antigenic map constructed with the antigenic distances thus predicted displays the same qualitative antigenic evolution patterns as those observed in the empirical maps, even across two antigenic cluster jumps (S2C Figure).

We assess the extent to which this algorithm could inform vaccine strain selection by providing phenotypic profiling of newly collected viral samples by evaluating its accuracy in predicting cluster transitions, and particularly the antigenic novelty of future vaccine strains. To determine whether newly sampled viruses are antigenically novel, we train the RFA up to a specific point in time and predict the antigenic distance between all antigen/serum pairs constituted by an element of the current antigenic cluster and one collected in the year following the reference time point. We thus predict the antigenic novelty of each new viral sequence given what is known till the year preceding that virus’ sampling. The mean antigenic distances predicted for each year of collection shows not only the expected peaks upon a cluster jump but also some degree of within cluster antigenic drift – Figure 2. More strikingly, the vaccine strains defining each cluster are quite consistently predicted to be antigenically novel (with the exceptions of EN72, and SI87) one year in advance. Whereas the prediction for EN72 is limited by the availability of HI data to inform the RFA, the BK79 to SI87 antigenic cluster transition is actually predicted to have occurred earlier (around 1985). The mean predicted antigenic novelty of viruses collected in 2002 is quite low, since most of those actually belong to the SY97 antigenic cluster. Nevertheless, the FU02 vaccine strain sequence is clearly predicted to be antigenically distant to previously circulating viruses and there is a substantive difference when comparing antigenic distance predictions for future members of the current circulating cluster compared to elements of a new yet unobserved cluster – S2 Table. Overall, the concordance of the blindly predicted antigenic novelty of H3 viruses with the empirical measurements is quite striking.

#### *Molecular evolution algorithm*

##### Investigating selection for differences in physiochemical properties.

We developed a simulation model that generates viral sequences with as many mutations as required from a given reference viral sequence, following Kimura’s nucleotide evolution model [25]. In practice, we generate a given number of nucleotide substitutions starting from a specific strain sequence, and record the resulting amino acid substitutions. These nucleotide substitutions are distributed randomly between the observed set of polymorphic amino acid sites in HA1, with nucleotides within each amino acid being randomly selected for mutation, whilst keeping to a 2:1 transition to transversion ratio. If a thus simulated substitution is synonymous, another one is generated at the same amino-acid position until we obtain a non-synonymous change. This algorithm was implemented in Matlab® R2016a.

By comparing the physiochemical properties of the simulated viral sequences thus generated against those collected in nature, we can evaluate influenza’s evolutionary plasticity, and infer the antigenic significance of the observed mutations in the context of all possible mutations. S4 Figure displays the resulting changes in polarity and hydrophobicity when imposing a range of mutations relative to the HK68 strain (using a 2:1 the transition to transversion ratio) randomly allocated to the polymorphic sites present in our alignment. Each box represents the variability in the differences in physiochemical properties of 100,000 thus simulated viruses relative to HK68.

##### Molecular evolutionary models

Here, we explore 3 molecular evolutionary models, expressing quite different mutation distribution profiles, all assuming a neutral model of DNA substitution with a fixed 2:1 transition to transversion rate ratio:

- Model 1: A 7-site model which is the most likely to display canalized antigenic evolution, in which mutations are restricted to the sites identified by Koel and colleagues [26] to be cluster determinants.
- Model 2: RFA 25-site model, where mutations are restricted to the 25 most significant amino acid positions as predicted by the random forest trained on the full dataset.
- Model 3: Generalized model described below.

Let us define a set of HA1 amino acid positions *ρ* consisting of a set of sorted (descending) RFA significant sites, a set of epitopes *ε* not included in *ρ*, and the (randomized) set of remaining polymorphic sites in the HA1 region as *π*. A truly general evolutionary model would allow mutations in *ρ, ε*, and *π*. It is challenging to parameterize such a model, given that we clearly have not observed all the evolutionary steps between any pair of viruses (particularly across antigenic clusters). We can, however, catalogue where mutations between all observed viruses have occurred and calculate the proportion that fits into each subset defined above (S6 Figure). The proportion of mutations falling on subset *ρ*, defined as the 25 most significant amino acids as determined by a RFA trained on the full dataset, is remarkably stable (~50%), with a slight peak in the 10-20 mutations range and obviously saturating when more than 25 mutations are observed. The proportion of mutations falling on subset *ε* is constantly around 15%, whereas the proportion of mutations on any epitope is always approximately 33% (not shown). We thus have a mutational profile that consistently follows a 4:1:2 distribution of mutations across the *ρ*, *ε* and *π* subsets.

#### *Simulating evolutionary trajectories*

Combining the predictive ability of the GP map with our viral sequence simulation tool, we can investigate the limits of plausible antigenic evolution. Given the mean antigenic distance between any two viruses belonging two adjacent antigenic clusters to be 4.46 (
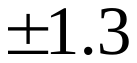
) on average [1], we simulate thousands of viral sequences from a specific ancestor and filter them according to their predicted antigenic distance to past clusters, according to two simple rules: 1) the simulated viruses must have predicted antigenic distance values within the 3-5 range relative to all viruses within the reference cluster; 2) predicted distances to viruses in all other past clusters must be greater than 6. Taking the SY97 cluster as reference and the SY97→FU02 transition as an example, we implement 12-18 mutations starting from strain NL/124/01 (4-6 mutations at each step, selecting one of the intermediate viruses as template for the next step), and filter the simulated viruses such that the final subset conforms with the antigenic distance rules mentioned above. The resulting antigenic map illustrates how closely the observed antigenic trajectory of H3 influenza can be represented by one single simulated lineage (with the same standing genetic diversity as the Fujian 02 cluster) obeying the mentioned antigenic rules. Indeed, simulated lineages with mutations either restricted to amino acid positions found to be significant in explaining the antigenic distances in the dataset comprised of viruses in FU02 and its two preceding clusters (S9A Figure), or to actual polymorphisms across SY97 and FU02 clusters (S9B Figure) display a striking overlap with the FU02 cluster (notice that predicted distances between FU02 viruses and simulated ones are also accounted for here).

An important question is whether the observed lineages are the only possible genetic evolutionary pathways for the virus to generate significant antigenic novelty. The linearity of the antigenic maps suggests strong immune-mediated selection for influenza viruses to maximize their antigenic distance to previous viruses [1,8]. Here we consider how selection for antigenic novelty shapes the possible evolutionary trajectories the virus might follow. We do so by considering three evolutionary processes: (1) antigenically unfiltered evolution where the simulated viruses are not parsed according to their antigenic novelty compared to current circulating ones; (2) filtered evolution with simulated viruses only being accepted as likely if certain predicted antigenic distance criteria are met, parsing out simulated viruses that are predicted to not be antigenically different from vaccine strain ones; (3) filtered step-wise evolution in which viruses are filtered much like in (2) only with the caveat of having intermediate bottlenecks, with a single simulated intermediate viruses serving as a seed for the next mutational step. We assume physio-chemical constraints are pervasive and thus mutations are bounded by the observed underlying limits of antigenic plasticity for each amino acid. In the 3-step mutational process we assign 5 to 8 mutations on the same SY97 ancestral strain mentioned above, on a first step, and impose the resulting predicted antigenic distance between simulated virus and ancestor to be greater than 2. The second and third mutational steps are in all similar to the first, except the additional mutations imposed on the seeding simulated viruses have to result in a predicted antigenic distance to the ancestor greater than 3. Antigenic distances to all other previous clusters is required to be over 6 throughout.

We assessed the validity of these evolutionary processes for each of the molecular evolutionary models, simulating 100,000 viruses for each process-model combination – S11 Figure. Each generated virus is evaluated in terms of the predicted antigenic distance (the RFA is trained only on strains isolated from SY97 and earlier clusters) to its ancestor and to all other simulated viruses (for process (3), only to the viruses resulting from the last mutational step).

Even in the filtered, step-wise evolutionary process there is substantial antigenic disparity across simulated viruses with the same number of mutations to their ancestor. It is then of relevance to investigate the predicted antigenic distances within and across simulated viral lineages. Out of all the simulated viruses meeting the imposed antigenic filter at the end of the second mutational step, two are selected to serve as seeds in the last mutational step, and antigenic maps are generated from the predicted antigenic distances using a set of 10 simulated viruses per simulated lineage as sera. In the main text we display the result of 10,000 simulated viruses per step, with each lineage being represented by viruses with 15-24 mutations to the ancestor. Several such simulations were conducted with identical results and we chose one of the most representative for illustration (Figures 4D, E of the main text).

### ***Simulating cluster jumps***

To further characterize the predictability of influenza’s antigenic evolution we try to re-create several cluster jumps. This involves selecting one evolved strain from each reference cluster and simulate sufficient genetic evolution that would normally result in a cluster jump, *i.e.*, imposing 12-18 amino acid mutations. We adopt the generalized model and impose all antigenic selection relevant effects (‘plasticity’, ‘filtering’, and ‘seeding’). The seeding effect is implemented as a 3-step mutational process as before. Here we aim to better characterize the predicted antigenic distances of the simulated viruses to the observed next antigenic cluster by keeping track of the RFA predicted antigenic distances to all members of that cluster. This information, together with the predicted distances to all other past observed clusters, allowed us to perform a *k-means* analysis to identify sufficiently distinct groups of simulated viruses (representing different antigenic evolutionary lineages). We found that the resulting optimal number of evolved ‘lineages’ was 3 for all simulated cluster jumps, suggesting there is a finite number of near-orthogonal evolutionary trajectories viruses can undertake in antigenic space. One of those trajectories was consistently antigenically very close to the observed next clusters. We then identified which specific mutations were the most significant for the differentiation of the simulated viruses into distinct antigenic groups by performing a discriminant analysis of principal components (DAPC) [27]. This type of analysis aims to summarize the genetic differentiation between groups, while minimizing within-group variation. The DAPC confirmed that the simulated lineages similar to the observed clusters shared critical mutations suggested to be determinant for cluster jumps [26]. These are summarized in S4 Table.

**S1 Table. Genbank references for viral sequences used in this study.**

| Antigen | Genbank reference | Antigen | Genbank reference |
| --- | --- | --- | --- |
| BI/15793/68 | 383213096 | **NL/179/93** | 49339268 |
| BI/16190/68 | 49339048 | **PA/287/93** | 49339234 |
| BI/16398/68 | 383213115 | **SG/6/93** | 49339282 |
| BI/808/69 | 383213153 | **YA/56/93** | 49339274 |
| BI/908/69 | 383276564 | **YA/61/93** | 49339276 |
| BI/17938/69 | 383276583 | **YA/62/93** | 49339278 |
| BI/93/70 | 383213172 | **NL/399/93** | 49339304 |
| BI/2668/70 | 383213191 | **SD/9/93** | 49339288 |
| BI/6449/71 | 383213210 | **LY/672/93** | 49339306 |
| BI/21438/71 | 383276602 | **LY/1803/93** | 49339308 |
| BI/21801/71 | 383276621 | **NL/357/93** | 49339294 |
| BI/6022/72 | 383213248 | **NL/371/93** | 49339298 |
| BI/21793/72 | 383213267 | **NL/372/93** | 49339296 |
| BI/23290/72 | 383276640 | **NL/398/93** | 49339302 |
| BI/23337/72 | 383213286 | **NL/440/93** | 49339300 |
| BI/552/73 | 383276678 | **OS/2219/93** | 49339316 |
| BI/748/73 | 383213304 | **OS/2352/93** | 49339318 |
| BI/3517/73 | 383276697 | **ST/20/93** | 49339292 |
| BI/5146/74 | 383213323 | **VI/104/93** | 49339284 |
| BI/5930/74 | 383276735 | **WE/59/93** | 49339286 |
| BI/5931/74 | 383276754 | **LY/1815/93** | 49339310 |
| BI/7398/74 | 383276773 | **LY/22686/93** | 49339312 |
| BI/9459/74 | 383276792 | **LY/23602/93** | 49339314 |
| BI/2600/75 | 383276849 | **SP/3/93** | 49339392 |
| BI/2813/75 | 383276868 | **HK/1/94** | 49339320 |
| BI/628/76 | 383276906 | **HK/2/94** | 383214241 |
| BI/1761/76 | 383276925 | **HK/55/94** | 49339378 |
| BI/2271/76 | 49338984 | **HK/56/94** | 49339326 |
| BI/5029/76 | 383276944 | **SA/15/94** | 49339322 |
| BI/5657/76 | 383276963 | **SA/25/94** | 49339324 |
| BI/6545/76 | 383276982 | **GD/25/93** | 2271380 |
| AM/1609/77 | 383277001 | **SL/142/93** | 383279076 |
| BI/3895/77 | 383277020 | **SL/160/93** | 383279095 |
| RD/5828/77 | 383277039 | **EN/7/94** | 383214203 |
| RD/8179/77 | 383277058 | **JO/33/94** | 49339330 |
| NL/209/80 | 383277096 | **FI/338/95** | 49339336 |
| RD/577/80 | 49339062 | **JO/47/94** | 49339328 |
| NL/233/82 | 383279874 | **GE/A9509/95** | 49339334 |
| NL/241/82 | 383277153 | **HK/32/95** | 49339340 |
| NL/330/85 | 383277229 | **GE/6447/91** | 49339124 |
| ST/10/85 | 49339074 | **LY/1189/91** | 49339170 |
| CO/2/86 | 49339076 | **LY/1276/91** | 49339176 |
| NL/450/88 | 49339078 | **LY/1337/91** | 49339178 |
| NL/620/89 | 49339104 | **LY/1373/91** | 49339174 |
| VI/2/90 | 49339116 | **LY/1594/91** | 49339172 |
| LY/1149/91 | 49339166 | **LY/23672/91** | 49339180 |
| MA/G102/93 | 49339258 | **LY/24103/91** | 49339182 |
| MA/G252/93 | 49339272 | **LY/24222/91** | 49339184 |
| NL/241/93 | 49339290 | **ST/20/91** | 49339208 |
| NL/1/95 | 49339332 | **FI/218/92** | 49339188 |
| WE/4/85 | 49339070 | **GE/5113/92** | 49339192 |
| SI/2/87 | 762858 | **HO/56798/92** | 49339218 |
| VI/7/87 | 2275516 | **HO/56829/92** | 49339214 |
| BI/4791/81 | 383213399 | **HO/56941/92** | 49339216 |
| BI/10684/82 | 383277134 | **OV/31/92** | 49339206 |
| OS/13676/83 | 383277191 | **NI/3126/92** | 407185927 |
| NL/333/85 | 383277248 | **NL/823/92** | 49339186 |
| BA/1/79 | 6578823 | **PA/320/92** | 49339134 |
| PH/2/82 | 7331124 | **PA/325/92** | 49339136 |
| CE/1/84 | 2275474 | **PA/407/92** | 49339138 |
| LE/360/86 | 2275546 | **PA/417/92** | 49339140 |
| CC/2/88 | 2275550 | **PA/424/92** | 49339142 |
| ST/12/88 | 49339082 | **PA/457/92** | 49339144 |
| EN/427/88 | 49339080 | **PA/490/92** | 49339148 |
| GE/5007/89 | 49339084 | **PA/512/92** | 49339150 |
| SH/11/87 | 348125 | **PA/548/92** | 49339152 |
| GU/54/89 | 2271054 | **PA/564/92** | 49339154 |
| HK/1/89 | 49339088 | **PA/597/92** | 49339158 |
| NL/650/89 | 49339106 | **RD/100540/92** | 49339194 |
| NL/738/89 | 49339028 | **ST/8/92** | 49339212 |
| SP/35/89 | 49339092 | **FI/339/95** | 49339338 |
| SP/40/89 | 49339096 | **HK/3/95** | 2271234 |
| WE/5/89 | 49339086 | **FI/381/95** | 49339362 |
| ME/2/90 | 49339108 | **HK/38/95** | 49339342 |
| SU/1/90 | 49339114 | **HK/49/95** | 49339344 |
| CA/1/91 | 49339122 | **HK/55/95** | 49339348 |
| EN/260/91 | 49339120 | **LY/2279/95** | 49339354 |
| EN/261/91 | 49339132 | **NA/933/95** | 2271174 |
| MA/G12/91 | 49339128 | **NL/271/95** | 49339352 |
| NL/816/91 | 49339126 | **VI/75/95** | 49339346 |
| AM/4112/92 | 49339200 | **WU/359/95** | 49339350 |
| ES/1285/92 | 49339162 | **BR/8/96** | 49339364 |
| FI/220/92 | 49339190 | **GE/3958/96** | 49339360 |
| MA/G58/92 | 49339204 | **HK/20/96** | 49339356 |
| NI/3129/92 | 49339198 | **HK/42/96** | 2271262 |
| NL/819/92 | 49339130 | **HK/357/96** | 2271146 |
| NL/935/92 | 49339164 | **HK/358/96** | 6552545 |
| PA/583/92 | 49339156 | **HK/434/96** | 6552513 |
| PA/614/92 | 49339160 | **NL/91/96** | 49339358 |
| SA/8/92 | 49339222 | **SP/1/96** | 49339368 |
| SA/23/92 | 49339224 | **HK/1/97** | 49339374 |
| SA/27/92 | 49339226 | **NE/491/97** | 49339390 |
| ST/7/92 | 49339210 | **AU/10/97** | 49339372 |
| VI/68/92 | 2271118 | **HK/280/97** | 49339376 |
| BE/353/89 | 49339102 | **NL/300/97** | 49339380 |
| LY/1182/91 | 49339168 | **SY/5/97** | 6552527 |
| BE/352/89 | 762860 | **NL/5/98** | 49339388 |
| PA/467/92 | 49339146 | **LY/1781/96** | 6318563 |
| TI/5957/92 | 49339202 | **JO/10/97** | 6552586 |
| AT/211/89 | 49339112 | **OS/21/97** | 14133839 |
| SP/34/89 | 49339090 | **OS/244/97** | 14133845 |
| SP/36/89 | 49339094 | **NL/414/98** | 49339382 |
| SP/53/89 | 49339098 | **NL/462/98** | 49339386 |
| VI/1/89 | 49339100 | **MW/10/99** | 49339008 |
| ME/5/90 | 49339110 | **NL/301/99** | 49339022 |
| SH/24/90 | 49339118 | **NL/3/00** | 49339012 |
| BE/32/92 | 49339230 | **NL/427/98** | 49339384 |
| FI/247/92 | 49339246 | **TE/1/77** | 18158213 |
| NL/938/92 | 49339220 | **GF/V728/85** | 49339072 |
| SE/C273/92 | 49339228 | **PM/2007/99** | 110825856 |
| ST/12/92 | 49339242 | **NL/118/01** | 49339014 |
| ST/13/92 | 49339244 | **NL/126/01** | 49339018 |
| UM/1982/92 | 49339238 | **NL/1/02** | 49339032 |
| UM/2000/92 | 49339240 | **NL/120/02** | 49339030 |
| AK/4/93 | 49339280 | **NL/18/94** | 49339010 |
| ES/5458/93 | 49339270 | **FU/411/02** | 81536028 |
| MA/G101/93 | 49339256 | **NL/22/03** | 49339038 |
| MA/G109/93 | 49339260 | **NL/213/03** | 49339040 |
| MA/G116/93 | 49339254 | **NL/217/03** | 49339042 |
| MA/G122/93 | 49339264 | **NL/222/03** | 49339044 |
| MA/G130/93 | 49339266 | **FI/170/03** | 49339034 |
| NL/3/93 | 49339232 | **NL/124/01** | 49339016 |
| NL/17/93 | 49339236 | **NL/20/03** | 49339036 |
| NL/101/93 | 49339248 | **HK/1/68** | 6470272 |
| NL/115/93 | 49339250 | **EN/42/72** | 6470274 |
| NL/126/93 | 49339252 | **PC/1/73** | 91125488 |
| NL/165/93 | 49339262 | **VI/3/75** | 60754 |
| Serum | **Genbank reference** | **Serum** | **Genbank reference** |
| HK/1/68 | 193805284 | **BE/32A/92** | 383278065 |
| EN/42/72 | 392341388 | **HK/34/90** | 2271040 |
| PC/1/73 | 392342083 | **BE/32B/92** | 2271348 |
| VI/3A/75 | 383276887 | **SD/9/93** | 2271364 |
| LE/360/86 | 392345721 | **OS/2352/93** | 383214621 |
| TE/1A/77 | 392329358 | **GD/25/93** | 2271380 |
| BI/21793/72 | 383213267 | **JO/33/94** | 2271272 |
| BI/5930/74 | 383276735 | **GE/A9509/95** | 383279285 |
| AM/1609/77 | 383277001 | **FI/338/95** | 383279228 |
| RD/577/80 | 383213380 | **NA/933/95** | 383214412 |
| NL/209/80 | 383277096 | **LY/2279/95** | 383279342 |
| NL/233/82 | 383279874 | **SP/1/96** | 383214507 |
| NL/241/82 | 383277153 | **BR/8/96** | 383279969 |
| NL/330/85 | 383277229 | **AU/10/97** | 383279532 |
| ST/10/85 | 383277267 | **SY/5B/97** | 383214640 |
| NL/450/88 | 383277381 | **NL/5/98** | 383279988 |
| VI/2/90 | 383277723 | **WU/359B/95** | 383214488 |
| LY/1149/91 | 383277818 | **SY/5A/97** | 226955221 |
| MA/G252/93 | 383278924 | **MW/10/99** | 392349364 |
| NL/1/95 | 383279361 | **SH/31/80** | 380835315 |
| NL/172/96 | 3885875 | **NL/1/02** | 383214773 |
| BA/1/79 | 392331726 | **PM/2007/99** | 383214716 |
| PH/2/82 | 392344267 | **NL/126/01** | 383279627 |
| WE/4/85 | 383277286 | **NL/118/01** | 383214735 |
| VI/7/87 | 2275516 | **VI/3C/75** | 383276887 |
| SI/2/87 | 392346364 | **NL/18/94** | 383279209 |
| TE/1B/77 | 392329358 | **FU/411/02** | 383214754 |
| BA/2/79 | 148340564 | **NL/22/03** | 383214811 |
| CO/2/86 | 383277305 | **NL/124/01** | 383279608 |
| SH/11/87 | 383277324 | **HK/107/71** | 383213229 |
| NL/620/89 | 383277457 | **PH/2V/82** | 299791410 |
| GU/54/89 | 2271054 | **BE/32V/92** | 194304840 |
| BE/353/89 | 383279931 | **SY/5V/97** | 226955221 |

**S2 Table. One year ahead predicted antigenic distances discretized by whether the newly sampled viruses are members of the current circulating cluster or turn out to belong to a new antigenic cluster.**

|  | **Predicted distances** | | **Standard deviations** | |  |
| --- | --- | --- | --- | --- | --- |
| **Prediction year** | **Within cluster** | **Across clusters** | **Within cluster** | **Across clusters** | **Vaccine strain** |
| 69 | 0.67 | NA | 0.58 | NA |  |
| 70 | 0.00 | NA | 0.00 | NA |  |
| 71 | 0.33 | NA | 0.58 | NA |  |
| 72 | 0.67 | 0.00 | 1.15 | NA | 0.00 |
| 73 | 3.00 | 4.00 | 0.00 | NA |  |
| 74 | 1.55 | NA | 1.44 | NA |  |
| 75 | 1.36 | 2.00 | 1.63 | 1.41 | 2.00 |
| 76 | 0.27 | 1.50 | 0.90 | 0.93 |  |
| 77 | 1.00 | 3.50 | 0.77 | 0.89 | 4.00 |
| 78 | 1.00 | NA | 0.77 | NA |  |
| 79 | 1.00 | 1.00 | 0.77 | 0.99 | 1.00 |
| 80 | 1.18 | 4.50 | 1.25 | 2.73 |  |
| 81 | 0.64 | NA | 0.92 | NA |  |
| 82 | 0.27 | NA | 0.90 | NA |  |
| 83 | 1.00 | NA | 1.18 | NA |  |
| 84 | 0.91 | NA | 1.22 | NA |  |
| 85 | 1.82 | NA | 3.39 | NA |  |
| 86 | 1.91 | NA | 1.68 | NA |  |
| 87 | 1.91 | 0.22 | 1.58 | 0.44 | 2.67 |
| 88 | 1.82 | 2.82 | 1.92 | 1.63 |  |
| 89 | 1.76 | 3.84 | 1.48 | 1.84 | 2.67 |
| 90 | 1.74 | 3.88 | 1.20 | 1.45 |  |
| 91 | 1.67 | 3.03 | 1.11 | 1.00 |  |
| 92 | 1.54 | 2.46 | 0.61 | 1.68 | 3.67 |
| 93 | 1.77 | 4.96 | 1.20 | 2.34 |  |
| 94 | 1.74 | 3.84 | 0.80 | 1.95 |  |
| 95 | 2.10 | 4.87 | 1.32 | 2.30 | 9.00 |
| 96 | 2.23 | 3.70 | 0.96 | 1.80 |  |
| 97 | 2.12 | 2.91 | 0.91 | 1.85 | 3.00 |
| 98 | 2.11 | 2.50 | 0.99 | 2.07 |  |
| 99 | 2.12 | 4.25 | 1.08 | 2.12 |  |
| 00 | 1.99 | 4.00 | 1.08 | 2.09 |  |
| 01 | 1.98 | NA | 1.12 | NA |  |
| 02 | 1.90 | 2.67 | 1.11 | 2.11 | 2.67 |
| 03 | 2.04 | 3.71 | 1.12 | 2.04 |  |

**S3 Table. RFA 25 most significant sites.** Amino acid positions with high interaction indices are in bold italic.

| HA1 position | Equivalent position in Koel *et al* | RFA significance |
| --- | --- | --- |
| 294 | 278 | 1.81E-02 |
| *172* | 156 | ***1.48E-02*** |
| 174 | 158 | 1.12E-02 |
| *175* | 159 | ***1.10E-02*** |
| 163 | 147 | 1.06E-02 |
| *171* | 155 | ***1.06E-02*** |
| 296 | 280 | 1.03E-02 |
| 161 | 145 | 1.00E-02 |
| 162 | 146 | 9.85E-03 |
| 295 | 279 | 9.66E-03 |
| 148 | 132 | 9.65E-03 |
| *151* | 135 | ***9.22E-03*** |
| 207 | 191 | 8.92E-03 |
| 208 | 192 | 8.61E-03 |
| 150 | 134 | 8.53E-03 |
| *209* | 193 | ***8.35E-03*** |
| 152 | 136 | 8.30E-03 |
| *292* | 276 | ***8.27E-03*** |
| 153 | 137 | 8.21E-03 |
| 101 | 85 | 7.86E-03 |
| *149* | 133 | ***7.69E-03*** |
| 170 | 154 | 7.04E-03 |
| *147* | 131 | ***6.63E-03*** |
| 160 | 144 | 6.61E-03 |
| *173* | 157 | ***6.52E-03*** |

**S4 Table. Antigenic lineage defining mutations.** The amino acids predicted here to be significant for the discrimination of antigenically distinct lineages that overlap with those mutations proposed to cause cluster jump by Koel *et al.* (27) are underlined.

| Cluster transition | *Ancestral strain* | *Significant amino acids in differentiating antigenic lineages* |
| --- | --- | --- |
| BE89→BE92 | HK/34/1990 | 78, 137, 145, 155, 156, 189, 193 |
| BE92→ WU95 | NL/18/1994 | 45, 54, 81, 129, 137, 138, 144, 155, 193, 219 |
| WU95→SY97 | BR/8/1996 | 31, 54, 79, 158, 262, 276 |
| SY97→FU02 | NL/124/01 | 53, 78, 122, 135, 156, 167, 188, 192, 199, 214, 226, 262, 275, 289 |

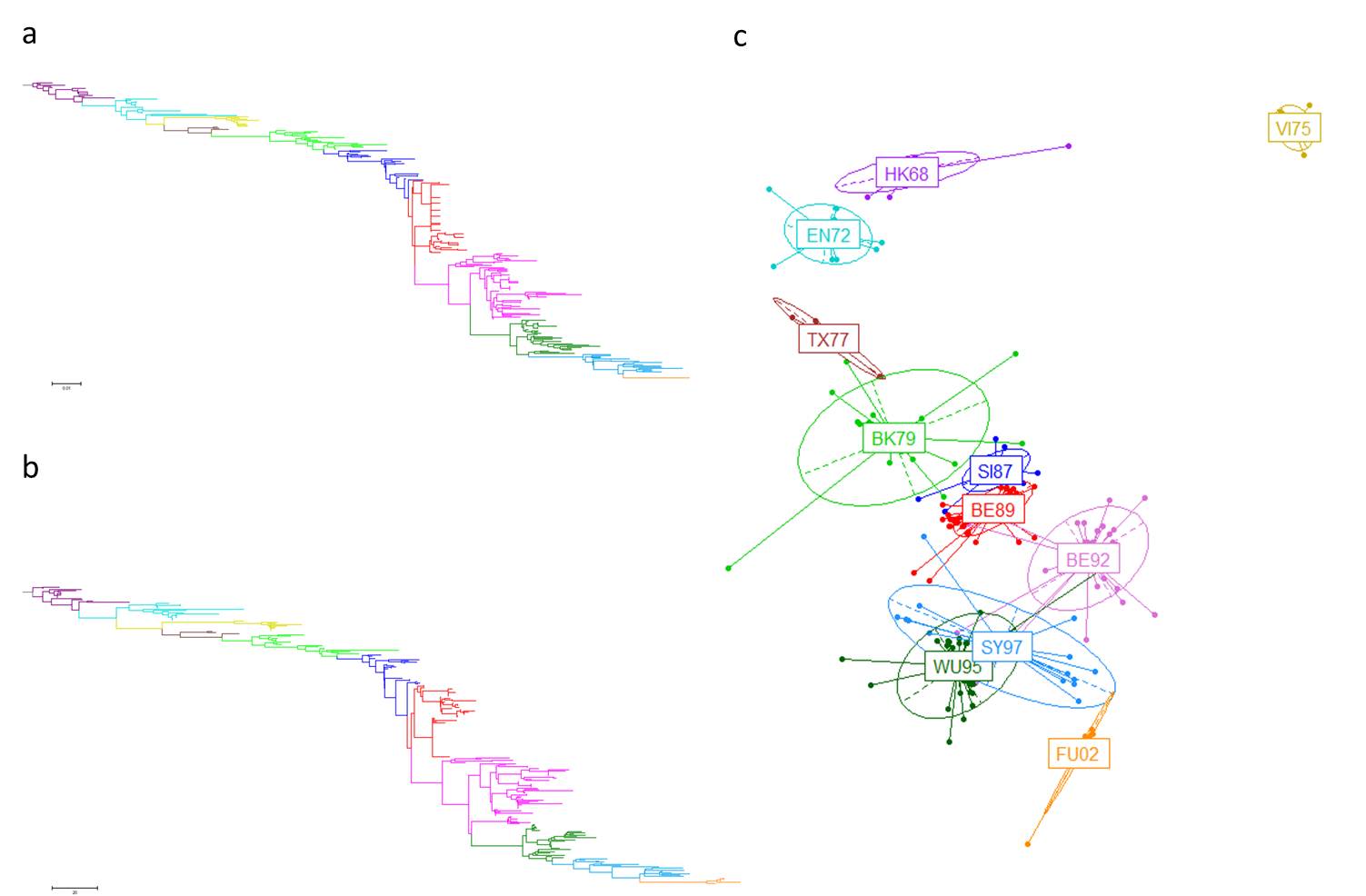

**S1 Figure**. **Correspondence between genetic distances and sum in polarity and hydrophobicity across the sequences.** (A) ML phylogenetic tree of all antigen sequences. (B) Dendrogram built from a distance matrix derived from the sum of differences in polarity and hydrophobicity in the antigen sequences. (C) MDS using the same matrix as in (B).

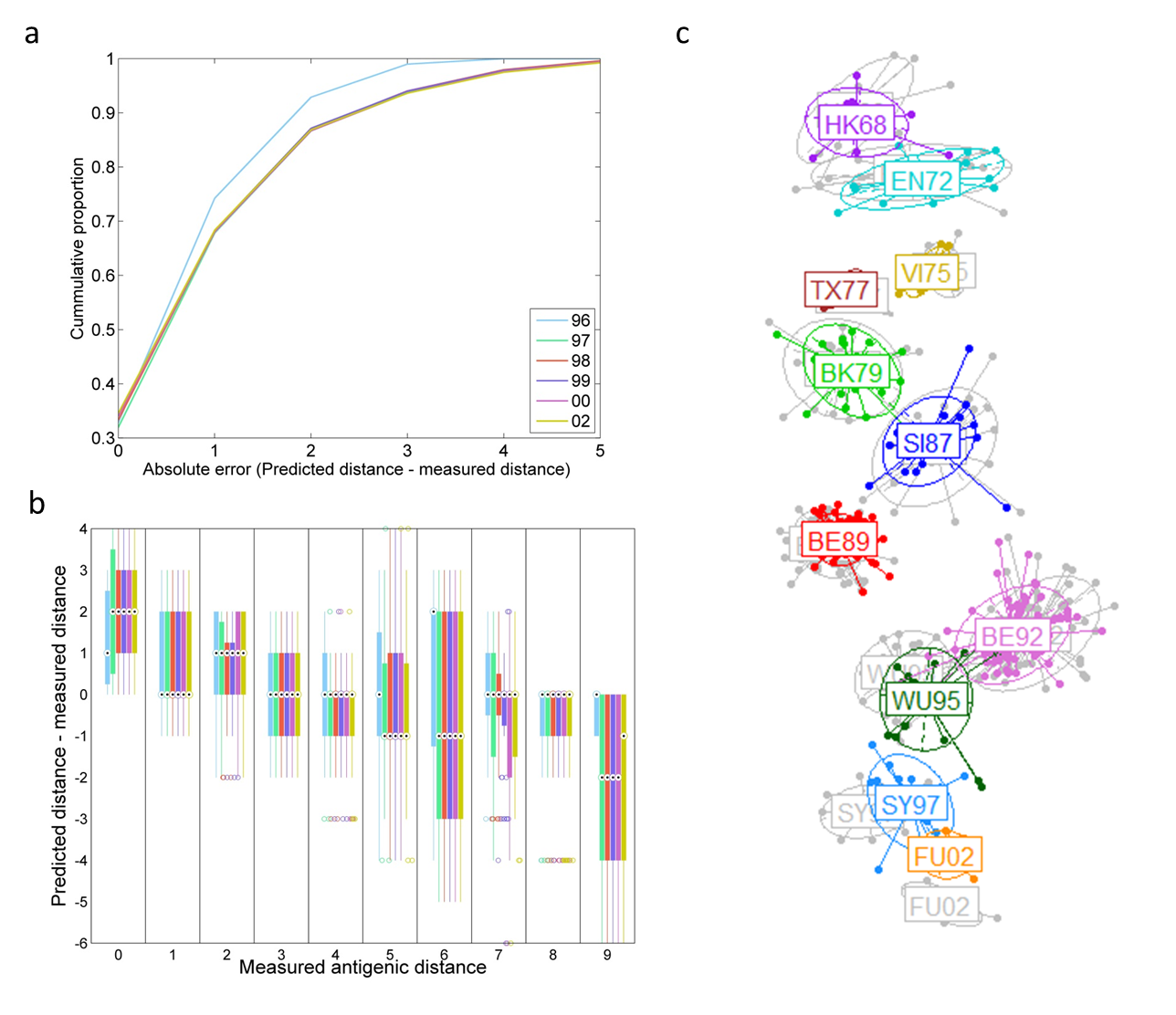

**S2 Figure**. **Prospective predictive power of the RFA algorithm with predictions made from a training set with data collected up to 1995.** (A) Cumulative curves representing the distribution of the prediction errors (on a log-2 scale) for the different prediction time windows. (B) Box plots of the prediction error for specific values of measured antigenic distance. For each box, the central mark represents the median, the edges mark the 25^th^ and 75^th^ percentiles, and the whiskers extend to the most extreme data points not considered outliers. Here the whiskers extend to +/–2.7*variance (corresponding to 99.3 coverage assuming the data are normally distributed). Each colour represents an incremental prediction time window as in (A). (C) Predicted antigenic map with predictions in colour overlaid on the map generated with full hindsight (in grey).

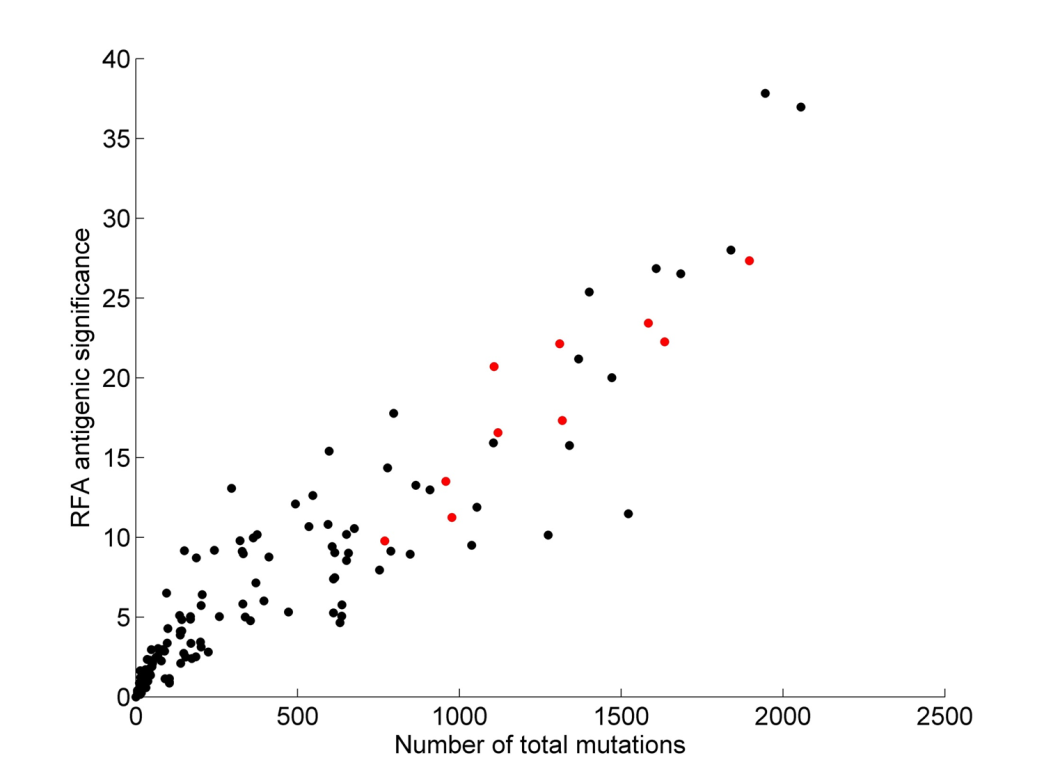

**S3 Figure.** **Correlation between the total number of mutations recorded at each position and the respective RFA (trained on the full dataset) significance.** Amino acid positions with high interaction indices are highlighted in red.

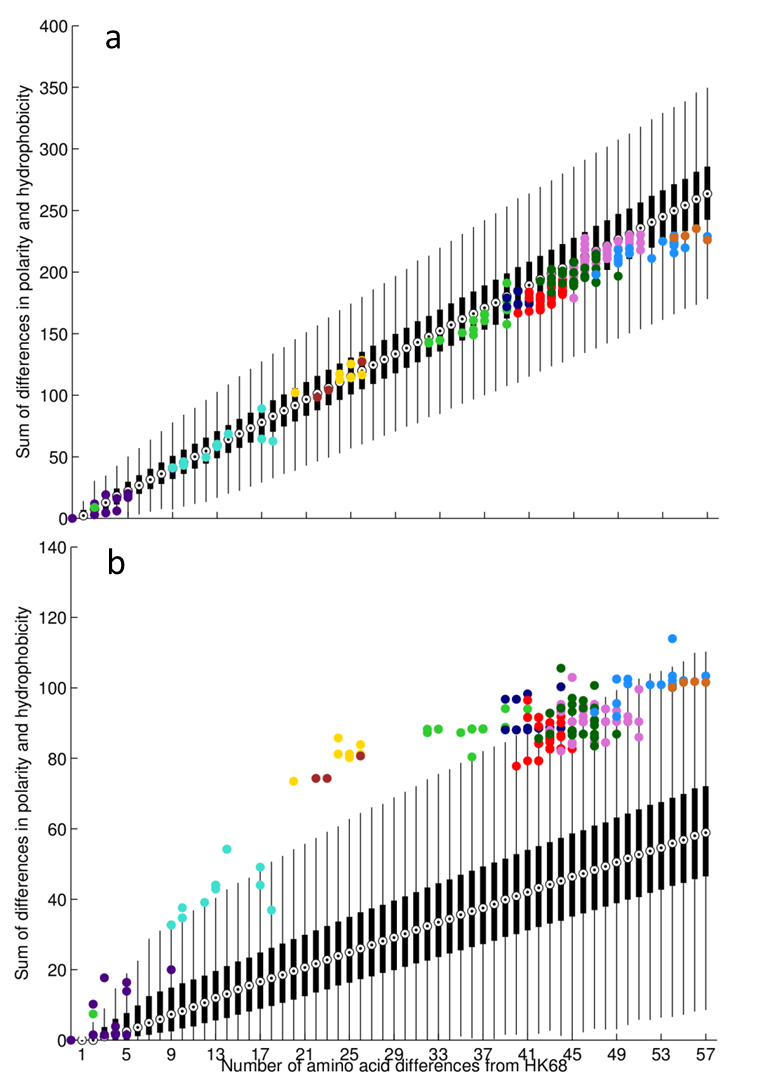

**S4 Figure**. **Evolution in HA1.** Each box plot in black show the variation in the sum of differences in amino-acid polarity and hydrophobicity between 100,000 viruses and their ancestral HK68 virus (given a specific number of mutations). The black circle in each box identifies the median, the edges mark the 25th and 75th percentiles, and the whiskers extend to the most extreme data points not considered outliers. (A) Shows differences across the full HA1 sequence, and (B) across epitope positions only. Isolated viruses are color-coded according to their assigned antigenic cluster.

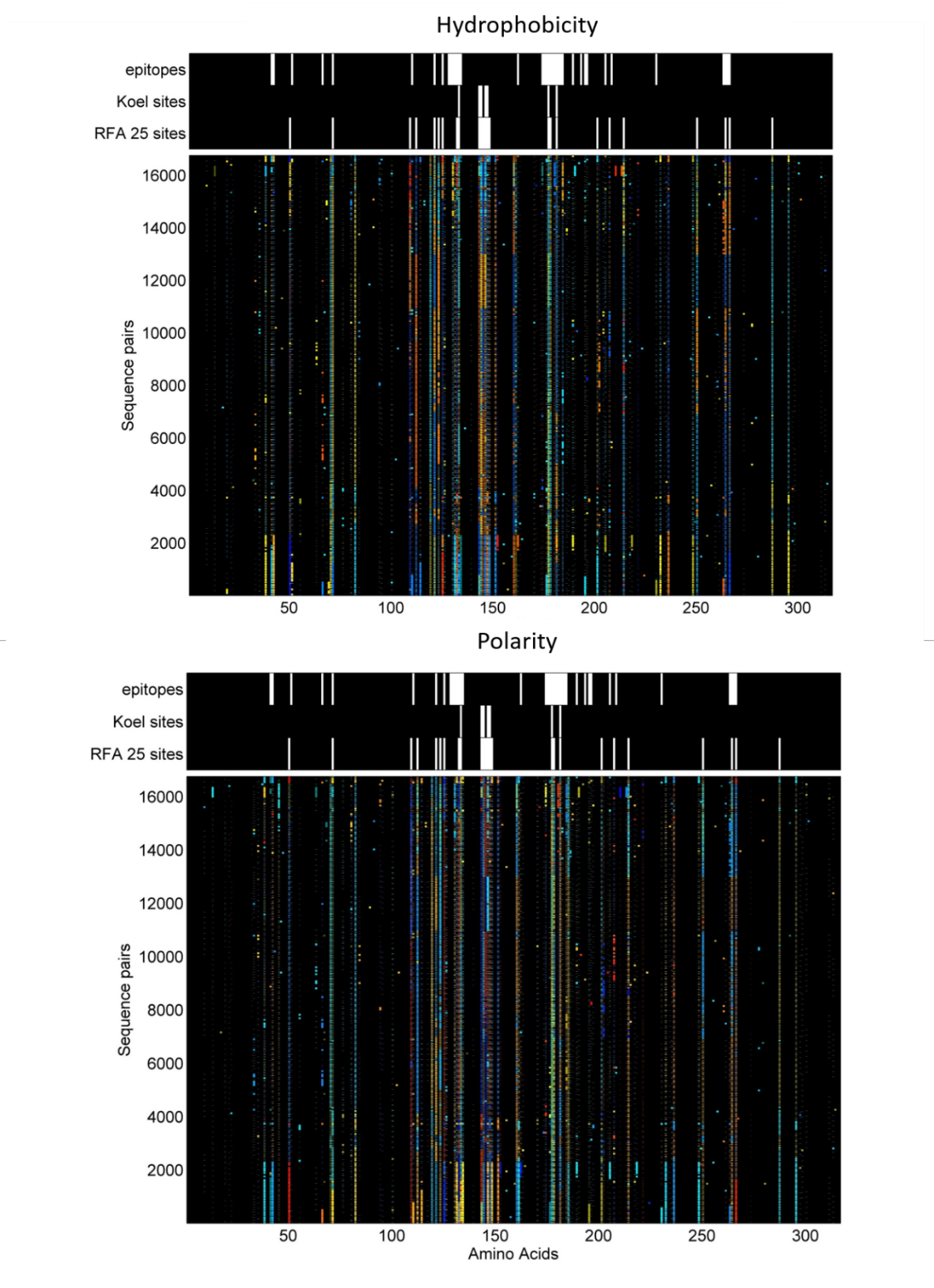

**S5 Figure**. **HA1 physiochemical plasticity**. Displays the observed difference in hydrophobicity (top) and polarity (bottom) for all antigen/serum pairs (rows) in each amino acid position (columns). Black indicates no difference, whereas blue and red represent the most extreme values (negative and positive, respectively).

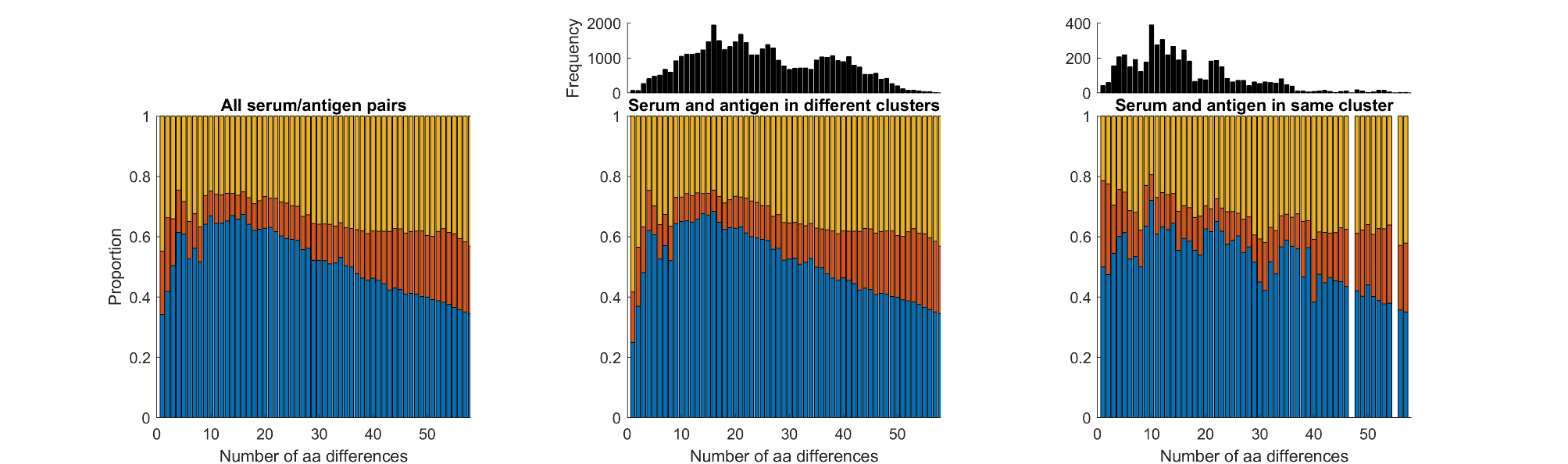

**S6 Figure**. **HA1 mutational profile**. Shows the distribution of mutations across three subsets of amino acids. Blue illustrates mutations that fall on the subset of RFA 25 most significant sites, in red we represent the known epitopes which are not included in the blue subset, and in orange the remaining polymorphic sites (not included in any of the other subsets).

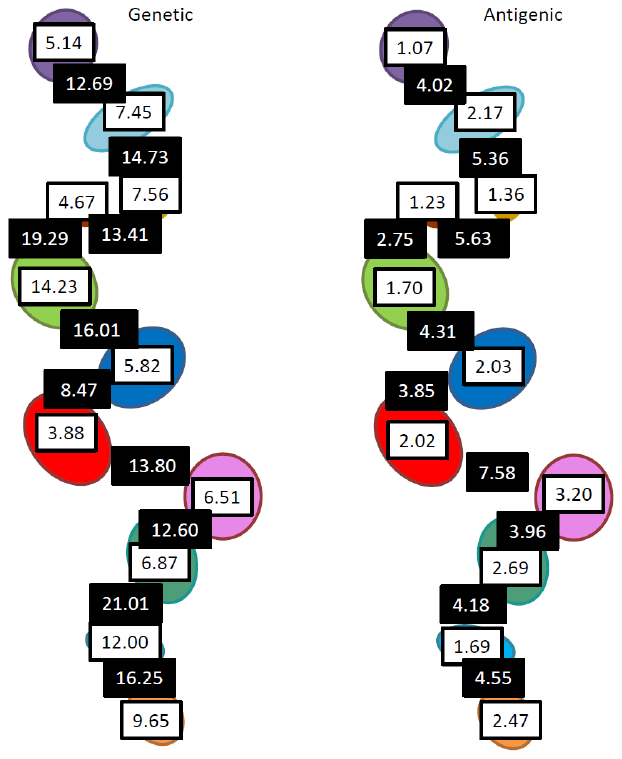

**S7 Figure**. **H3 Antigenic and Genetic Evolutionary patterns**. This sketch illustrates mean differences between members of the same cluster (white background boxes) and between viruses belonging to contiguous clusters (black background). The mean antigenic differences shown here are calculated from the HI titers measured in the considered antigen/sera pairs. The mean genetic differences are merely the mean number of amino acid differences between sequences of the same cluster or across contiguous clusters.

**S8 Figure**

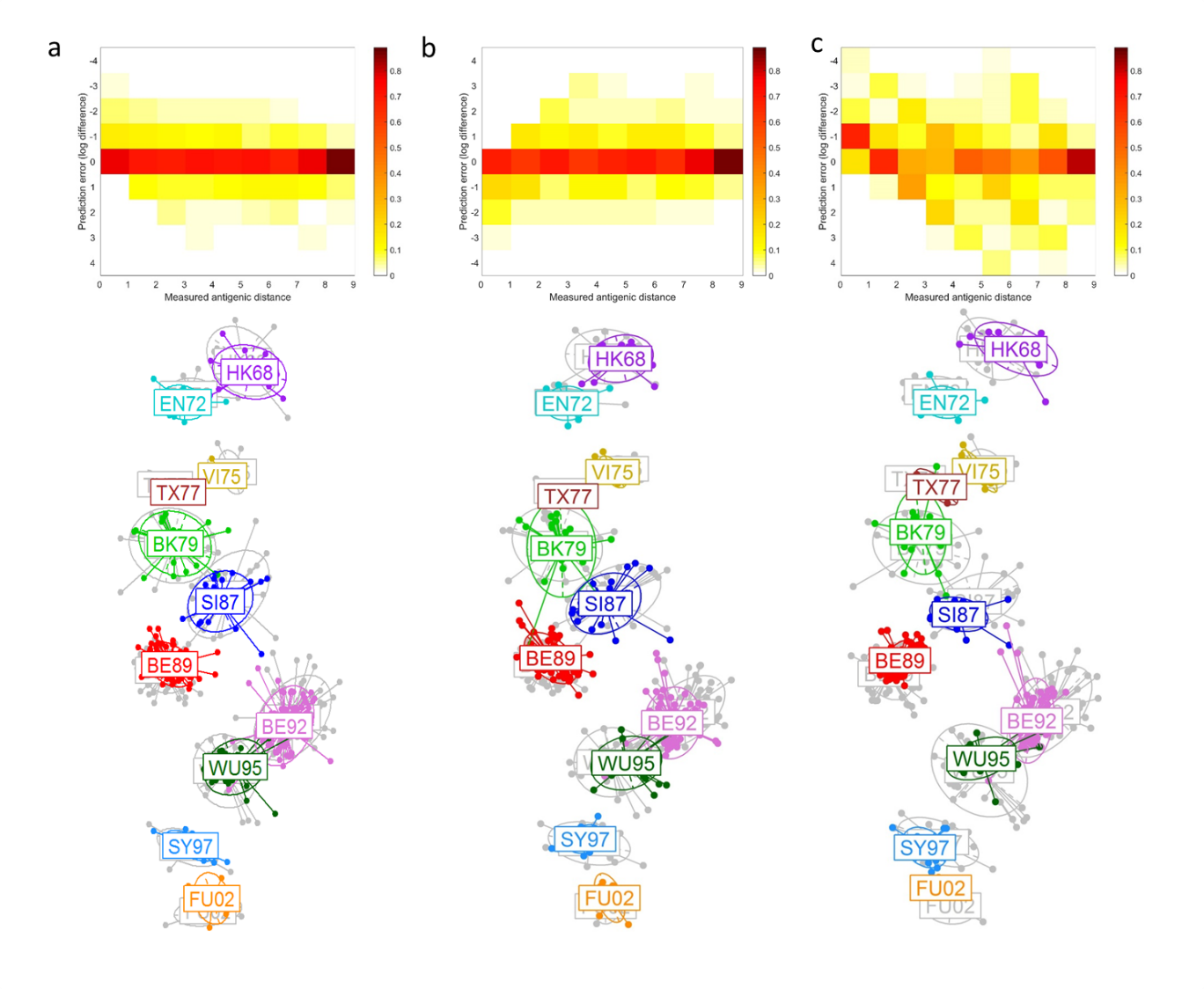

**S8 Figure**. **Evolutionary model accuracy**. Classification accuracy of the RFA looking at hydrophobicity and polarity differences in each of the considered evolutionary models: the generalized (A); the RFA 25 most significant sites (B) or the 7 sites model proposed by Koel *et al* (C). Superimposed 2-dimensional antigenic maps generated from the predicted antigenic distances (in colour) - when training the RFA on the respective subsets of amino acids - and from the HI titre data (in grey). The top panels give the prediction errors for each of the observed antigenic distance bins.

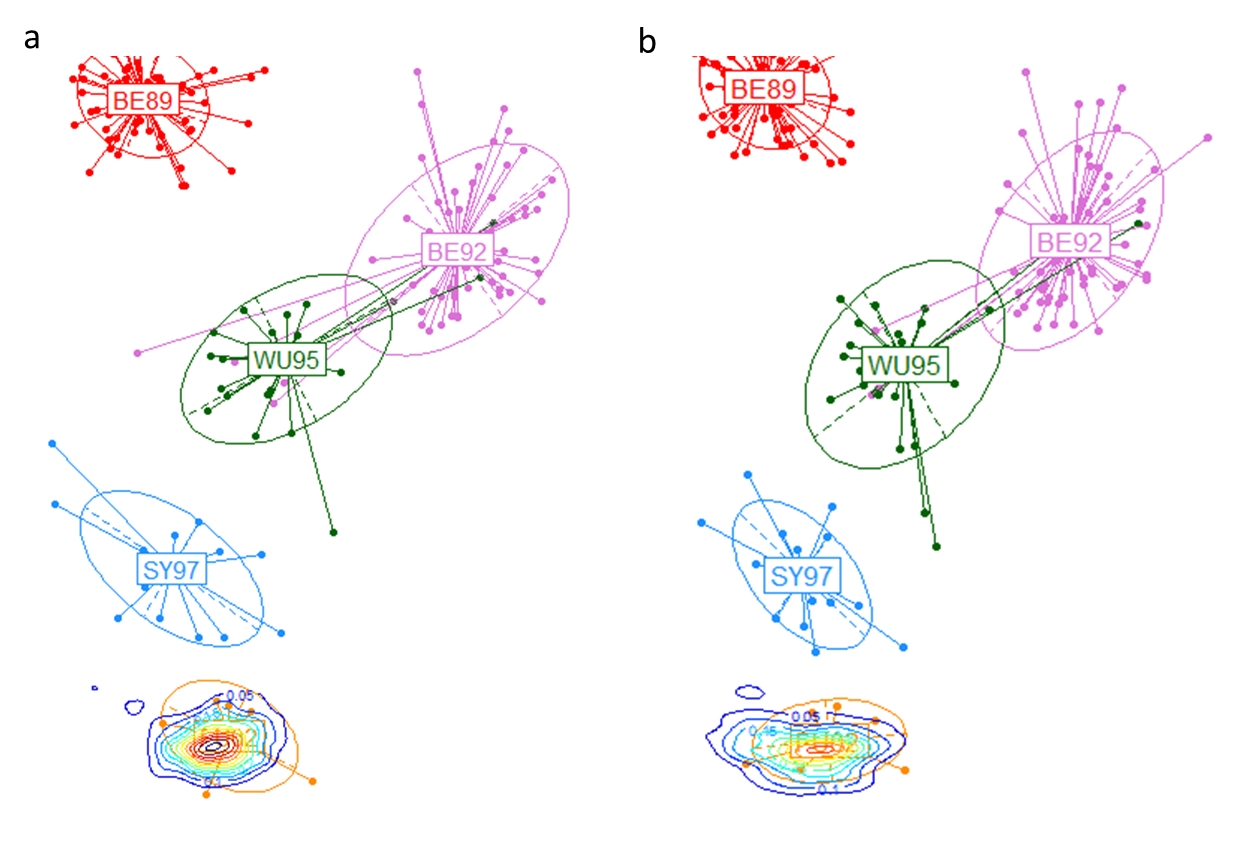

**S9 Figure**. **Antigenic maps of influenza’s predicted antigenic trajectories.** The contour lines represent the density probability that simulated viruses will fall on the respective delineated antigenic space. The simulated viruses were generated from mutations in: (A) significant positions to explain antigenic differences between clusters BE92 to FU02; (B) the polymorphic sites across the SY97and FU02 clusters. Each simulated virus has a total of 12-18 amino acid mutations relative to its ancestor.

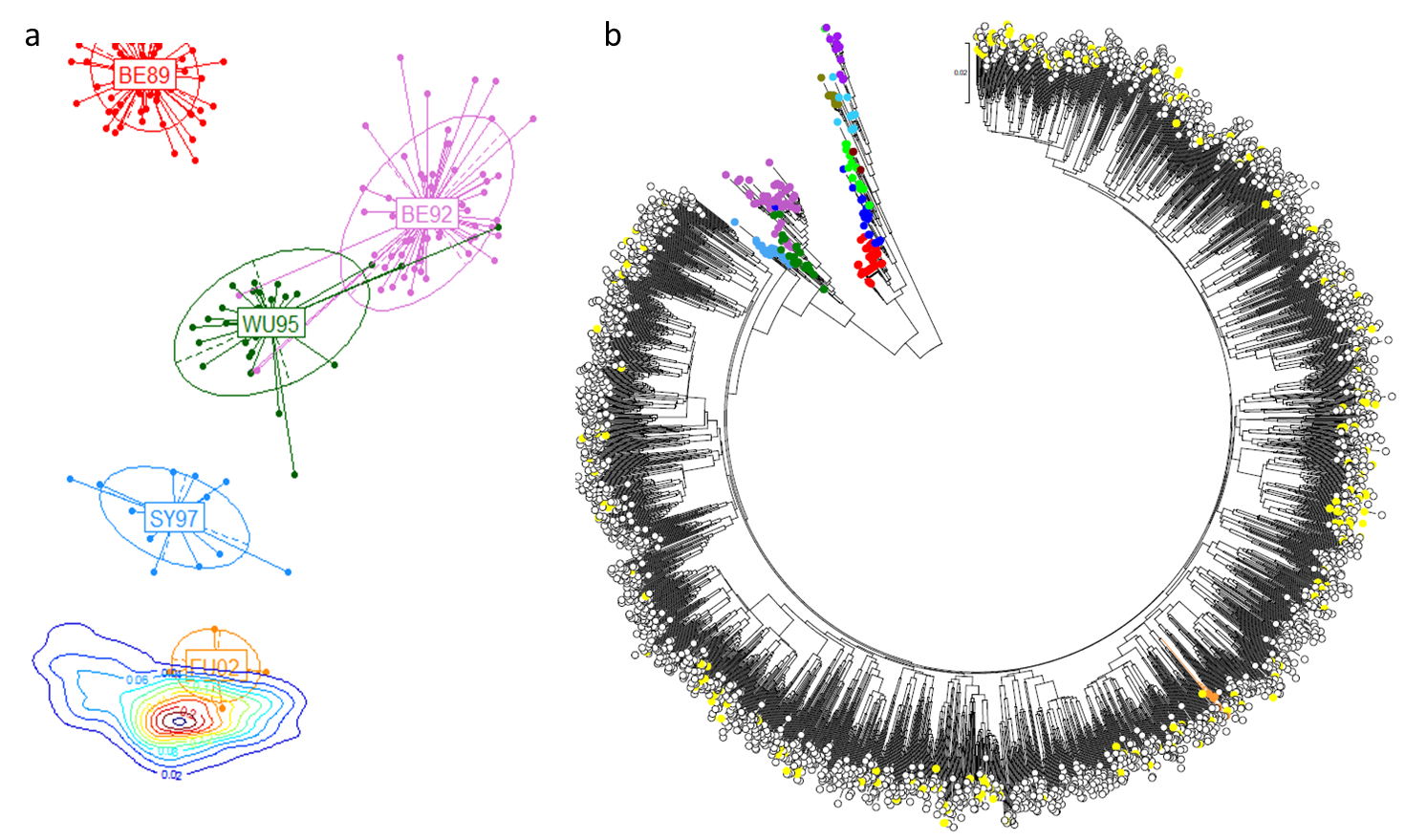

**S10 Figure. Antigenic trajectory unpredictability.** Antigenic map of influenza’s predicted antigenic trajectory using the generalized model (A). We implemented a mutational process similar to that in S9 Figure and impose an antigenic filter that only accepts simulated viruses with predicted antigenic distances to the ancestral viral cluster in [3, 5] and of over 6 to all preceding clusters. The contour lines represent the density probability that simulated viruses will fall on the respective delineated antigenic space. A maximum likelihood neighbor joining tree shows the genetic relationship between simulated viruses that did not meet the antigenic filter requirements (white), those that did (yellow), and the empirical viruses (colored according to their antigenic cluster as in all other plots). Note the empirical FU02 cluster in orange amidst the simulated viruses.

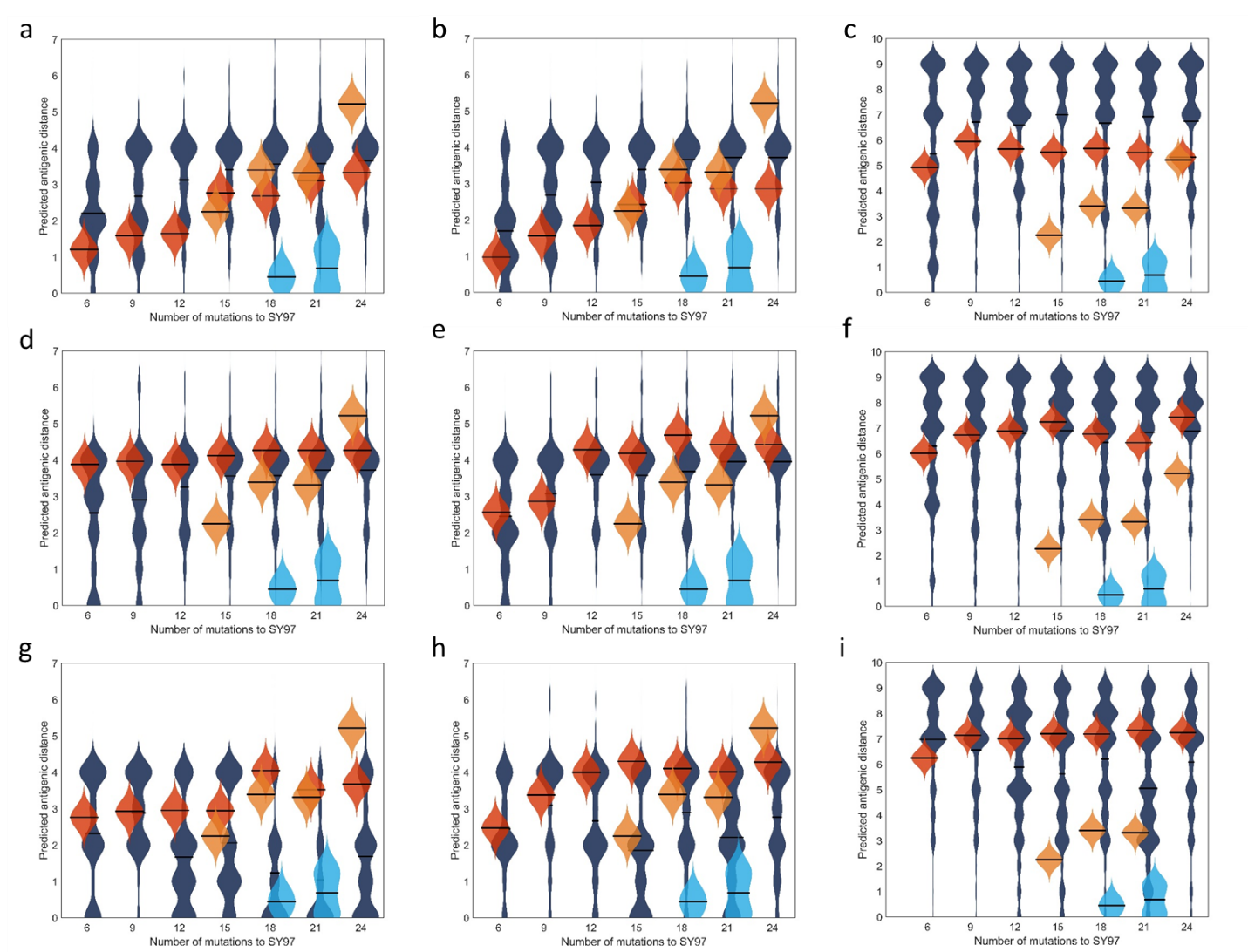

**S11 Figure.** **Antigenic trajectories and underlying evolutionary processes.** Unfiltered (top row); filtered (middle row); and filtered step-wise (bottom row) antigenic evolution from a specific ancestral strain under the 3 evolutionary models. The results for the generalized, 25-site, and 7-site models are depicted left to right respectively. The dark blue violin plots represent the distribution of antigenic distances (with the respective means as horizontal lines) between simulated viruses with X amino acid mutations relative to a virus in SY97. The red violin plots give the predicted antigenic distances across simulated viruses. Light blue violin plots give the empirical antigenic distances between viruses in SY97 and FU02, whilst the orange plots refer to the measured antigenic distances within the FU02 cluster.

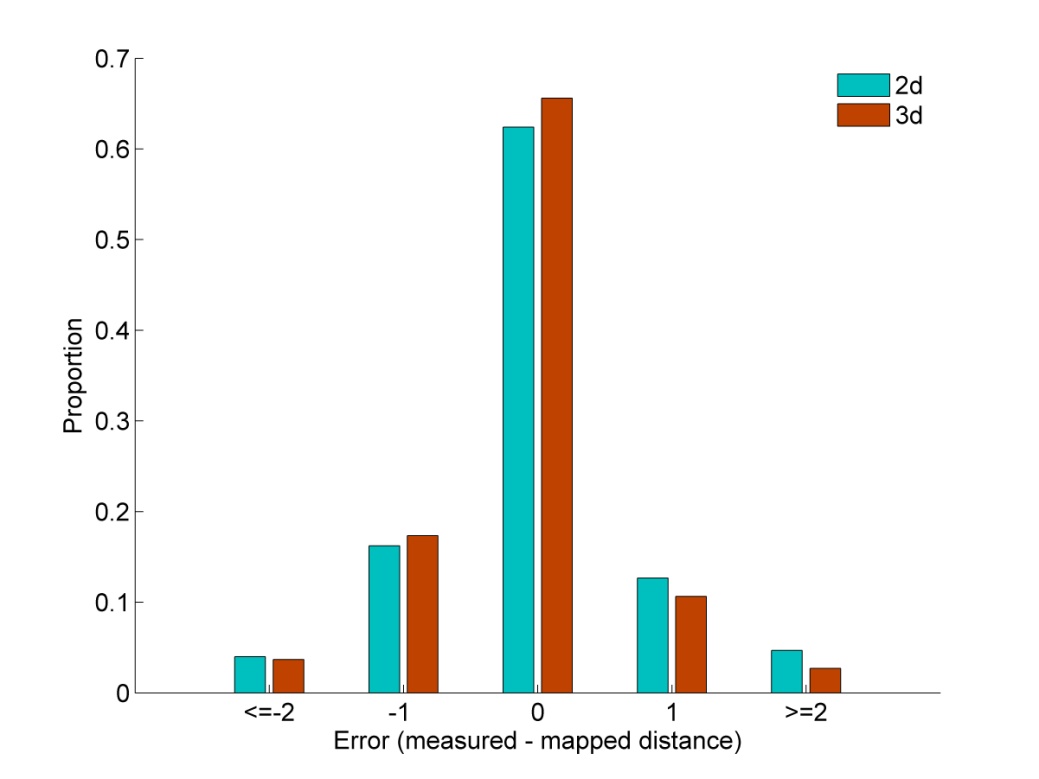

**S12 Figure**. **Antigenic map precision on different embeddings**. Comparison of the MDS error for the 2D and 3D representations of an antigenic map with two simulated lineages.

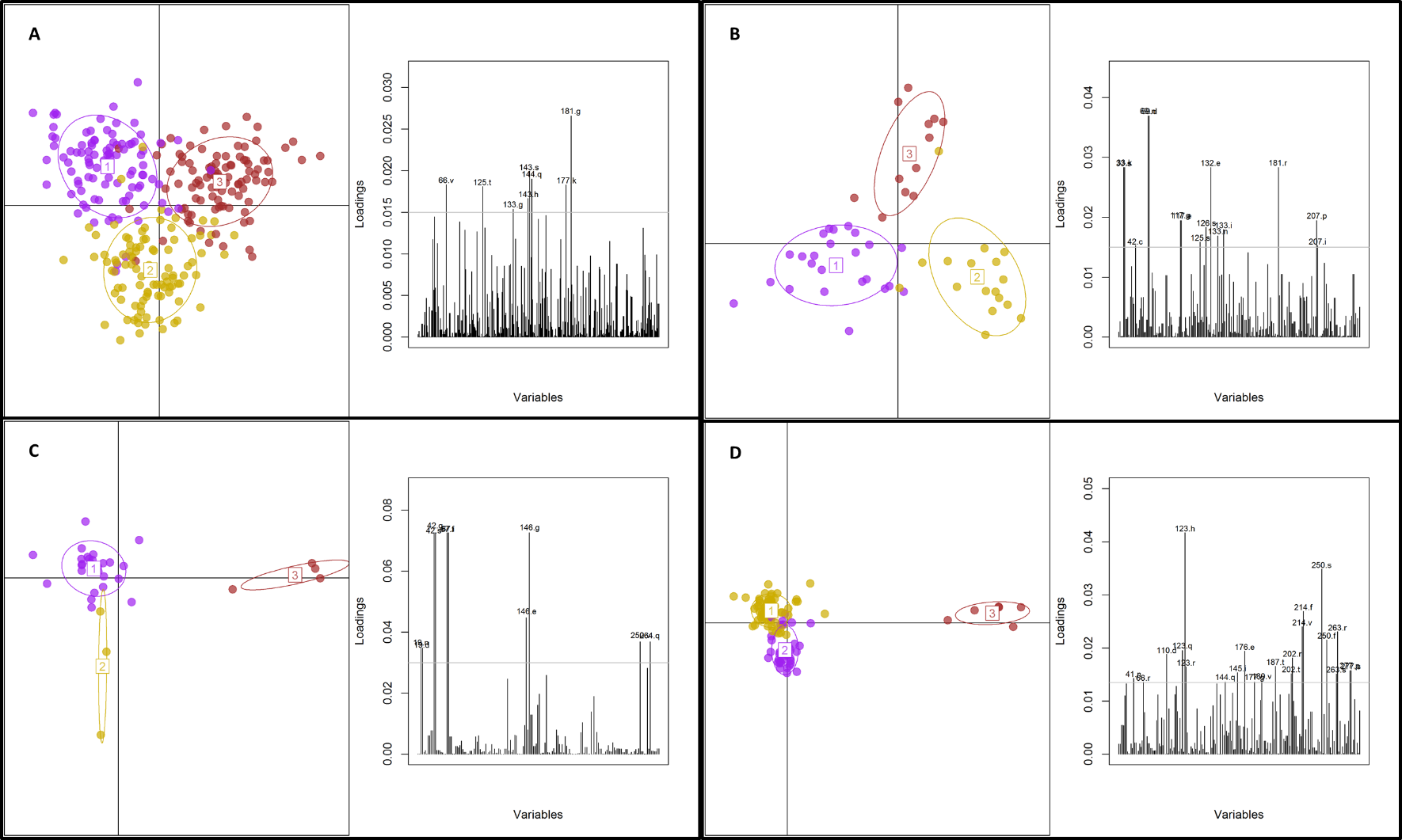

**S13 Figure**. **Genetic determinants of simulated viral lineages.** Each panel relates to the associated cluster jump in Figure 5 of the main text. The left plot of each panel shows the DAPC resulting discrimination of the separate groups given the genetic information. The contribution of each allele for the segregation of the viruses into groups is shown on the plots to the right. The dashed line serves only as an arbitrary threshold above which the individual allele labels are shown.

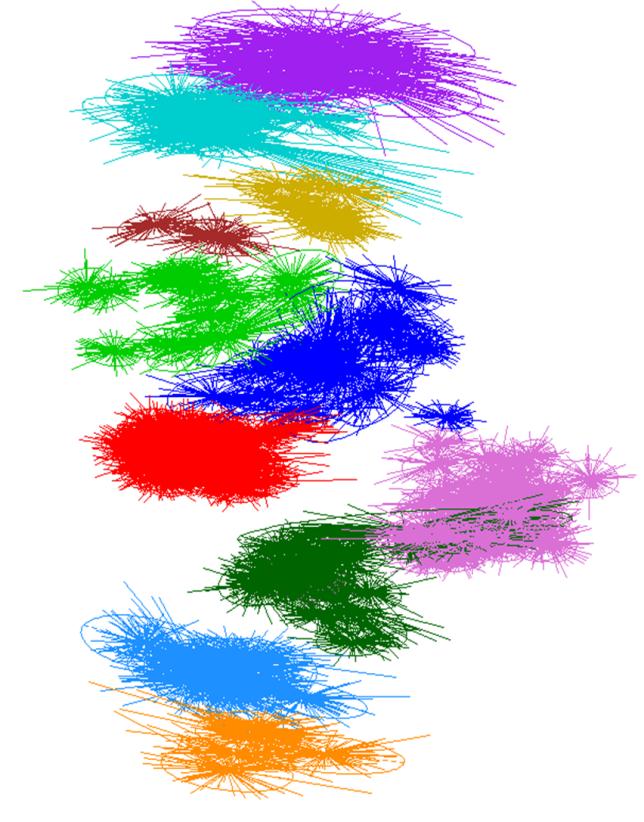

**S14 Figure**. **Uncertainty of the antigenic mapping algorithm of historic H3 influenza.** Each ellipsis represents the dispersal in the final proposed coordinates for each antigen (over 100 bootstrap runs of the mapping algorithm). Colors match the cluster assignments in.

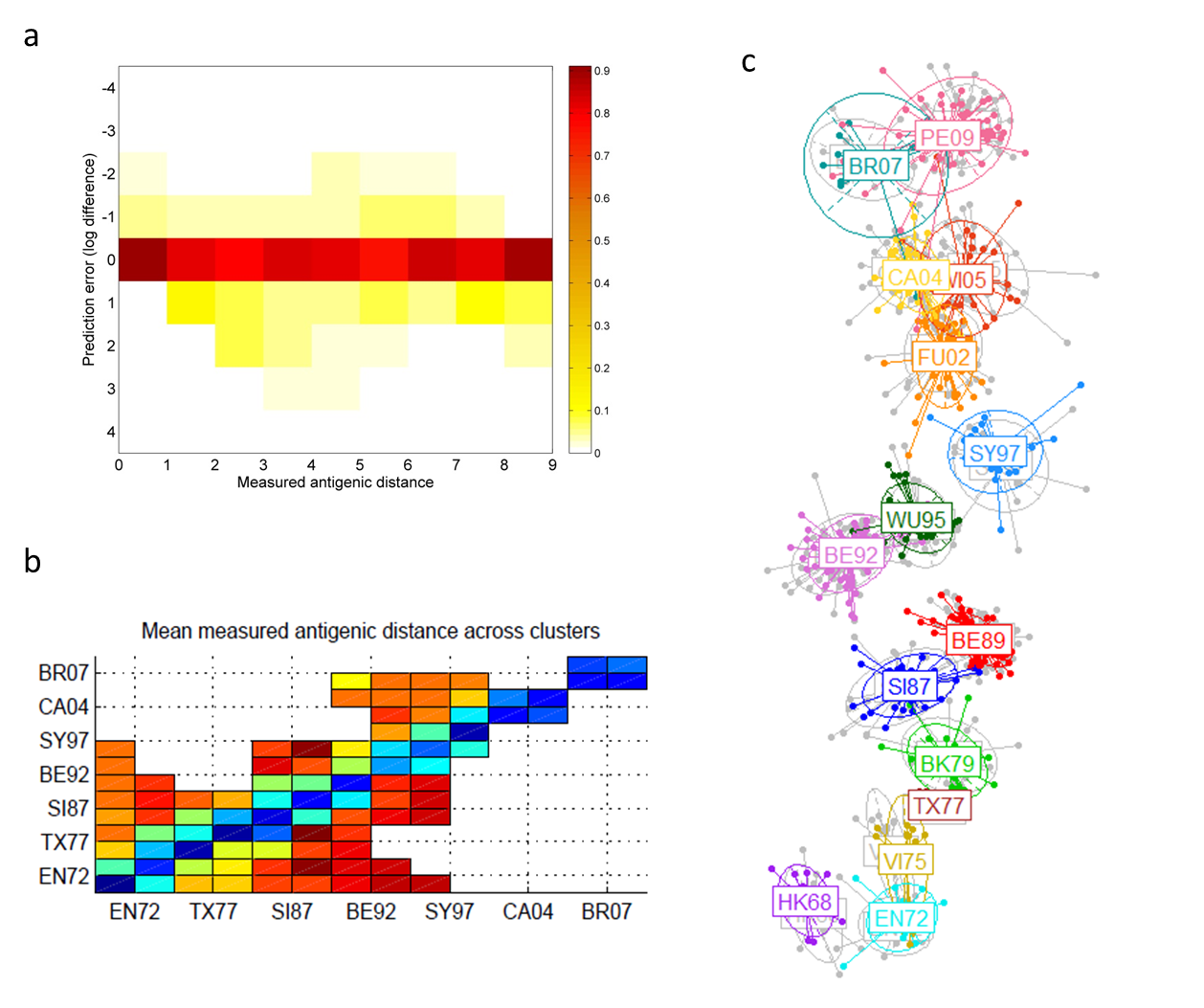

**S15 Figure.** **Classification accuracy of the RFA using hydrophobicity and polarity as antigenic proxy variables for an extended dataset**. (A) Surface plot showing the error in prediction of antigenic distance (distance being measured on a log-2 scale) for every antigen-serum pair. Antigen-serum pairs are grouped by measured antigenic distance on the horizontal axis. The colour of each rectangular pixel shows the proportion of pairs with at a particular measured antigenic distance with a particular prediction error. (B) Mean measure antigenic distance between antigen/serum pairs composed of elements of different antigenic clusters. The sparseness of this matrix of measurements increases significantly for more recent data. (C) Superimposed 2-dimensional antigenic maps generated from the RFA predicted antigenic distances (in colour) and from the HI titre data (in grey).
